# A dynamic cell recruitment process drives growth of the Drosophila wing by overscaling the Vestigial expression pattern

**DOI:** 10.1101/688796

**Authors:** Luis Manuel Muñoz-Nava, Hugo Ariel Alvarez, Marycruz Flores-Flores, Osvaldo Chara, Marcos Nahmad

## Abstract

Organs mainly attain their size by cell growth and proliferation, but sometimes also grow through recruitment of undifferentiated cells. Here we investigate the participation of cell recruitment in establishing the pattern of Vestigial (Vg), the product of the wing selector gene in Drosophila. We find that the Vg pattern overscales along the dorsal-ventral (DV) axis of the wing imaginal disc, *i.e*., it expands faster than the DV length of the pouch. The overscaling of the Vg pattern cannot be explained by differential proliferation, apoptosis, or oriented-cell divisions, but can be recapitulated by a mathematical model that explicitly considers cell recruitment. By impairing cell recruitment genetically, we find that the Vg pattern almost perfectly scales and adult wings are approximately 20% smaller. Furthermore, using fluorescent reporter tools, we provide direct evidence that cell recruitment takes place in a specific time between early and mid third-instar larval development. Altogether, our work quantitatively shows when, how, and by how much cell recruitment shapes the Vg pattern and drives growth of the Drosophila wing.

## Introduction

Organ growth during development is orchestrated by morphogens, which are signaling molecules that determine gene expression in a non-cell autonomous manner and also act as mitogens (Day and Lawrence, 2000; Schwank and Basler, 2010; Dekanty and Milán, 2011; Lander, 2011; Wartlick *et al*., 2011; Bryant and Gardiner, 2016; Vollmer *et al.,* 2017). However, growth may also be driven independently of morphogen-induced cell proliferation, by a growth-by-patterning mechanism in which a differentiation pattern expands at the expense of the incorporation of undifferentiated cells. This mechanism, also known as inductive assimilation or cell recruitment, has been reported to work in both vertebrate and invertebrate development such as in the eye (Heberlein *et al.,* 1995; Strutt *et al.,* 1995) and wing discs (Baena-López and García-Bellido, 2003; Zecca and Struhl, 2007a) of the fruit fly, *Drosophila melanogaster,* as well as in the mammalian thyroid (Fagman *et al.,* 2006) and kidney (Lindström *et al.,* 2018). Little is known, however, about the dynamic properties of patterning by cell recruitment and how much organ growth can be gained through cell recruitment relative to cell growth and proliferation. These questions are generally difficult to address during normal development because the dynamics of patterning and cell proliferation occur simultaneously and it is challenging to isolate the contribution of each of these mechanisms to organ size. The Drosophila wing imaginal disc is a useful system to tackle this problem since previous studies have partly identified the molecular players of recruitment (Zecca and Struhl, 2007a; Zecca and Struhl, 2010) and genetic tools allow manipulation and quantitative assessment of patterning and growth (Hariharan and Bilder, 2006; Beira and Paro, 2016).

In Drosophila, wing fate is determined by the expression of the wing selector gene, *vestigial* (*vg*), in the pouch of the wing imaginal disc (Williams *et al.,* 1991). The absence of *vg* results in loss of wing blade identity (Williams *et al.,* 1991; Williams *et al.,* 1993), whereas ectopic expression of *vg* in other imaginal discs could induce transformation into wing-like tissue (Williams *et al.,* 1994; Kim *et al.,* 1996; Halder *et al.,* 1998; Baena-López and García-Bellido, 2003). Therefore, adult wing size depends on the size and shape of the Vg pattern established during the larval stage.

*vg* expression is controlled at least by two enhancers, the *vg* boundary enhancer (*vg*^*BE*^) that responds to Notch signaling and results in high expression of Vg at the dorsal-ventral (DV) boundary (Irvine and Vogt, 1997), and the *vg* quadrant enhancer (*vg*^*QE*^) that requires Wingless (Wg) signaling (Kim *et al.,* 1996; Zecca and Struhl, 2007b) resulting in a gradient of Vg along the DV axis (Baena-López and García-Bellido, 2006). The lineage of cells abutting the DV boundary can maintain their transcriptional expression of *vg* through Polycomb/Trithorax Responsive Elements (Pérez *et al.,* 2011). The *vg*^*QE*^ is the responsive element of a cell recruitment process that depends on a feed-forward signal sent by Vg-expressing cells to non-expressing cells (Zecca and Struhl, 2007a). In 2010, Zecca and Struhl identified the following molecular details of the recruitment process (Zecca and Struhl, 2010): (1) Vg-expressing cells transcriptionally repress the protocadherin *dachsous* (*ds*), resulting in complementary patterns of Vg and Ds expression; (2) the boundary of the Vg/Ds expression domains facilitates Fat-Ds polarization, and (3) the polarization signaling induces nuclear shuttling of Yorkie (Yki), the transcriptional factor downstream of the Warts-Hippo tumor suppressor pathway, where it promotes *vg* expression transcriptionally. This process results in a new Vg-expressing cell, thus propagating the Vg pattern. As new cells are recruited into the Vg domain, the Ds pattern is pushed outwards radially by Vg-dependent transcriptional repression from the center of the wing pouch.

While previous studies have provided experimental evidence of the recruitment process by showing that Vg-expressing mosaics can induce expression of a *vg*^*QE*^ reporter non-cell-autonomously (Zecca and Struhl, 2007a; Zecca and Struhl, 2010), the contribution of cell recruitment to wild-type patterning and growth of the wing disc has not yet been investigated. Here, we precisely address this question by quantitatively examining the temporal dynamics of the Vg gradient and asking if these can be explained by a cell recruitment mechanism. We show that cell recruitment affects the shape of the Vg pattern and contributes to final size of the adult Drosophila wing.

## Results

### The Vg pattern overscales with respect to disc size along the DV axis

In order to investigate how the Vg pattern changes as a function of tissue size, we quantified Vg expression in wild-type wing discs during the third larval instar (85-120 h After Egg Laying (AEL) at 25°C) as a function of DV position in a central region of the pouch delimited by the dorsal and ventral epithelial folds (Fig. 1A; Fig. S1; Supplementary Material). These folds robustly appear in the disc at about 85 h AEL at 25°C (Sui *et al.,* 2012) and serve as references to define the length of the DV axis within the pouch area (Fig. 1A). To determine how the Vg pattern changes in discs of different sizes, we subdivided the discs into 4 groups according to their DV length, which is correlated with disc age (Fig. 1B, Fig. S2). Representative discs of each group stained for Vg and DAPI are shown in Figure 1C-F. Since fluorescence levels drop close to the folds due to tissue geometry (Fig. 1C’’- F’’), we used normalized DAPI levels to correct Vg expression (see Supplementary Information). After DAPI correction, Vg is expressed in a concentration gradient with maximum levels around the DV border and decreasing towards the ventral and dorsal folds in all groups (Fig. 1C’’’- F’’’). However, when we plotted the average of the normalized Vg patterns of each group in relative units (*i.e.*, % of DV length), we observed clear differences in Vg patterns among the groups (Fig. 1G). First, we noticed that the relative width at half maximum of the Vg gradient (width to DV length ratio) on average increases when plotted vs. DV length (Fig. 1H, Fig. S3). Note that this result is independent of how the discs were grouped and indicates that the Vg gradient overscales with respect to disc size, *i.e.*, that the Vg pattern expands further than what would be expected by uniform growth of the disc along the DV axis. Second, the Vg gradient also experiences shape changes between groups; particularly, we noticed that the slopes at the tails of the Vg gradient between groups 3 and 4 decrease, suggesting that Vg levels significantly increase in cells located at the edges of the wing pouch during the late third instar (Fig. S4). Finally, we noticed a ventral shift of the relative position of Vg maximum between groups 2 and 3 (Fig. 1I); this shift suggests that the dorsal compartment grows more than the ventral compartment during this time. In summary, we found that during normal development, the scale, shape, and symmetry of the Vg gradient changes with respect to disc size along the DV axis.

**Figure 1.**
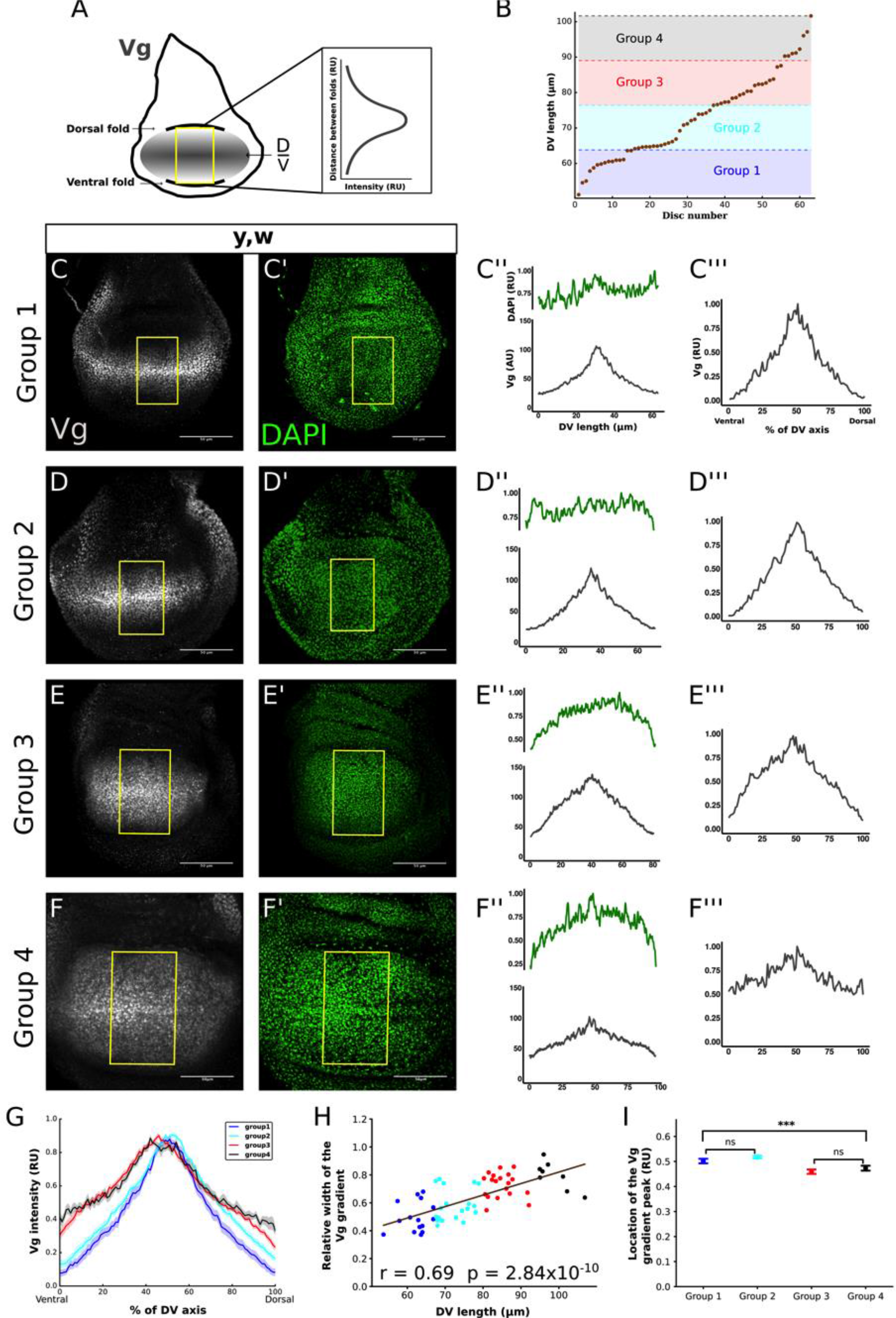
Spatio-temporal quantification of Vg along the DV axis in third-instar wing discs reveals overscaling of the Vg pattern to size. (A) Cartoon depicting the region within the wing pouch where the quantification of the Vg pattern was performed (yellow rectangle). This rectangular region is centered in the anterior posterior border of the disc and is delimited by the ventral and dorsal folds that separate the pouch from the hinge (throughout the paper, we use the distance between these folds as a measure of disc size). The intensity of Vg in each vertical line of pixels within the rectangle is averaged to obtain a spatial Vg expression along the DV axis. (B) Third-instar (ages 85-120 h After Egg Laying (AEL) at 25°C) yellow-white (*y*, *w*) discs (throughout the study, we refer to *y*, *w* discs as wild type) are classified into four groups according to the distance between dorsal and ventral folds (DV length). The groups are defined by dividing the shortest to the longest DV length into four intervals of equal length. (C-F) Representative images of discs within each group immunostained with a Vg antibody (C-F) and DAPI (C’-F’) and the quantification of the Vg patterns in the region defined in A in absolute (C’’-F’’) and relative (C’’’- F’’’) units (AU and RU, respectively). (In C’’-F’’, the quantification of DAPI patterns (green curves in the plots on the top of each panel) in RU are shown to indicate that disc geometry could affect the intensity levels close to the folds, especially in Group 3 and Group 4 discs). The quantifications of the Vg patterns in C’’’-F’’’ are normalized to DAPI levels (see Supplemental Information and Fig. S1). (G) Mean of the normalized Vg profiles (dark line) and Standard Error of the Mean (SEM, shaded area) from all the discs in each group (n=15, 21, 19 and 8 for Groups 1, 2, 3, and 4, respectively). (H) Relative width of the Vg gradient (defined as the width of the Vg pattern in RU at 0.5 divided by DV length) as a function of DV length; each dot in the graph corresponds to a different disc and is color-coded as in G. The solid line shows the linear regression of the data and the Pearson Correlation Coefficient, r, of the regression is displayed. The p-value corresponds to a Student-t statistical test assuming a zero-slope of the regression line as a null hypothesis. (Note that a positive slope corresponds to overscaling; a zero-slope, *i.e.*, when the null hypothesis cannot be rejected, corresponds to perfect scaling; whereas a negative slope corresponds to underscaling). (I) Comparison of the location of the Vg gradient peak in RU for the discs in each group; error bars correspond to the SEM. (Statistical significance is analyzed using a Kruskal-Wallis non-parametric test; ***, p-value<10-5. Pairwise statistical comparisons between groups 1 and 2, and groups 3 and 4 using a Mann-Whitney non-parametric test reveal non-statistical differences between these groups; ns, p-value<0.01).

### The dynamics of the Vg pattern is not a result of differential cell proliferation or apoptosis

We considered if overscaling of the Vg pattern could be explained by differences in cell proliferation or apoptosis between Vg-expressing *vs.* non-Vg expressing cells within the pouch. For example, cells near the edge of the pouch may be experiencing higher apoptosis rates or Vg-expressing cells may proliferate faster than cells outside of the Vg domain, explaining the overscaling of the Vg pattern. To test these possibilities, we first examined the expression of the pro-apototic marker Caspase 3 (Cas 3) throughout development and found that apoptosis occurs at low frequency and seems to be homogeneous in the wing pouch (Fig. S5); this is consistent with a previous study (Milán *et al.*, 1997). Second, we examined cell proliferation and found that during most of the third instar cell proliferation within the pouch is approximately uniform, except for cells at the DV boundary, confirming the results of previous studies (Fig. S6A; Schwank *et al.,* 2011; Wartlick *et al.,* 2011; Mao *et al.*, 2013). We conclude that the overscaling of the Vg pattern cannot be accounted by differences in cell proliferation or apoptosis between cells within and outside the Vg domain.

### Mathematical modeling of cell recruitment predicts the overscaling of the Vg pattern

In order to investigate if the dynamics of the Vg gradient could be explained by a cell recruitment mechanism, we modeled the distribution of Vg in the wing pouch by means of a multi-scale model (see Supplementary Material for full description). The model combines an ordinary differential equation to account for the rate of change of Vg concentration in each cell (*Vg*^*i*^) with a 2D Cellular Potts Model (CPM; Graner and Glazier, 1992; Glazier and Graner, 1993) describing the cellular dynamics (Fig. 2A and Fig. S7A, B). We assumed that cells produce Vg by two mechanisms, while there is only one sink modeled as a linear degradation. The first production term assumes that Vg expression responds to the concentration of a given non-scaling morphogen M (Fig. 2A and Supplementary Material). The second cellular source of Vg is our quantitative formulation of the recruitment mechanism. In particular, we proposed that this production term depends on the difference in Vg concentration between the actual cell and the average of the concentration of its neighbors by a second order Hill function (Fig. 2A and Fig. S7C). Hence, this production term becomes relevant when the concentration of the cell is different from the one of its neighbors and negligible when they are similar. As a control, we also considered a model without the recruitment term. To account for disc growth, we explicitly included homogeneous cell proliferation in which the cell cycle parameters were fitted to the average number of cells in each of the four groups defined in Fig. 1B (Fig. S8 and Supplementary Material). Schematics of the simulated Vg expression patterns corresponding to each of the groups are shown in Fig. 2B-E (without cell recruitment) and 2B’-E’ (with cell recruitment).

**Figure 2.**
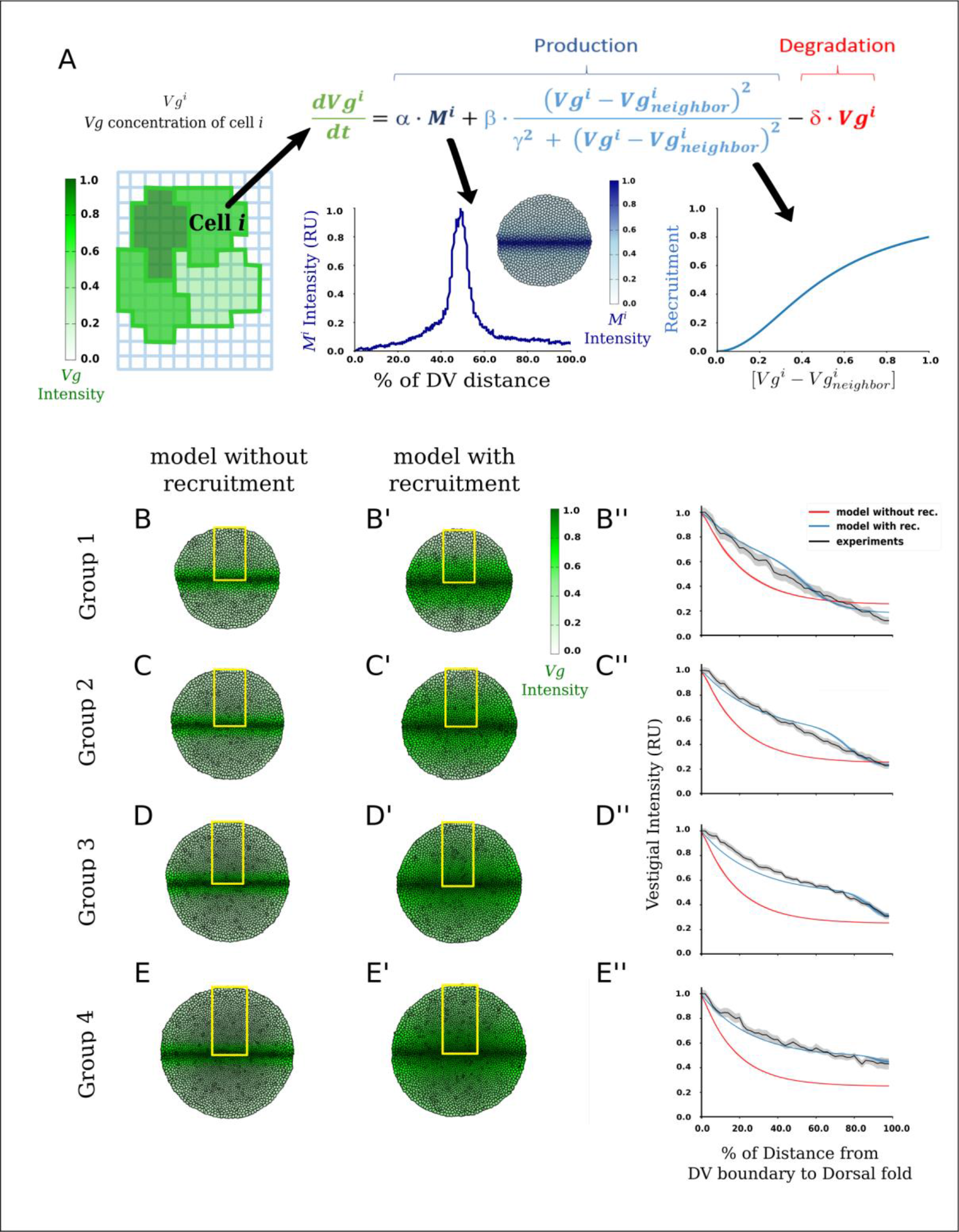
Multi-scale mathematical modeling with recruitment recapitulates Vg overscaling. (A) Outline of the model (see text and Supplemental Material for further details). The dynamics of Vg concentration in each cell is given by an ordinary differential equation including: a spatial-dependent production (first term) that models the concentration-dependent effect of source signal or morphogen M, our formulation of the recruitment mechanism (second term) that depends on a Hill function of the difference in Vg concentration between the actual cell and the average of its neighbors, and linear degradation (third term). As a control, we considered a model without the recruitment term. (B-E) Simulated tissues showing Vg profiles of the model without the recruitment term (B-E), and with the recruitment term (B’-E’) at four different simulated times corresponding to the average pouch areas of discs in Groups 1-4. (B’’-E’’) Simulated patterns of Vg in the dorsal compartment (obtained by horizontally averaging the simulated Vg patterns within the yellow rectangles in B-E and B’-E’) from the model with (blue curves) or without (red curves) the recruitment term, using parameter values that best fit the average experimental Vg profiles (black curves) in Groups 1-4.

We examined whether the models (with and without the recruitment term) could explain the spatiotemporal expansion of Vg shown in Fig. 1 by a fitting procedure where we varied the parameter values of each of the models and minimized a target function of the residuals between experimental and simulated Vg expression data. (For simplicity, we did not take into account differences between dorsal and ventral compartments, so we only compared experimental and simulated data in the dorsal compartment; Fig. S9). The best-fit simulation of the model with recruitment reasonably recapitulates the experimental overscaling in each of the four groups (Fig. 2B’’-E’’, blue line *vs*. black line; Fig. S10; Video S1; Supplemental Material) and the minimum of the target function is clearly identified within the parameter space (Fig. S11A-D, blue dots, minimum marked with a green circle). In contrast, the best-fit solution of the model without the recruitment term does not reproduce the experimental data (Fig. 2B’’-E’’, blue line *vs*. black line; Video S1).

Finally, since cell divisions in the wing pouch are oriented along the proximal-distal axis (González-Gaitán *et al.* 1994; Resino *et al.* 2002; Baena-López *et al.* 2005; Mao *et al.* 2013), we used our mathematical model to ask whether the overscaling effect could be enhanced by assuming that cell divisions are oriented radially. However, essentially the same fit was achieved in a model that assumes radially-oriented cell divisions vs. a model where cell divisions are randomly oriented (Fig. S12). We conclude that a mathematical model encoding a mechanism of cell recruitment may explain the overscaling dynamics of the Vg gradient regardless of the orientation of cell divisions.

### Blocking Vg expression in ds-expressing cells disrupts the overscaling of the Vg gradient and nearly results in perfect scaling

To verify the predictions of our mathematical model, we expressed an interference RNA for *vg* (*vg*^*RNAi*^) under *ds* control (*ds*>*vg*^*RNAi*^) using the Drosophila Gal4-UAS system (Brand and Perrimon, 1993; Fig. S13A). Since cell recruitment works by expanding Vg expression at the expense of *ds*-expressing cells (Zecca and Struhl, 2010), we expect cell recruitment will be blocked in *ds*>*vg*^*RNAi*^ discs. Moreover, if cell recruitment is indeed responsible for the overscaling of the Vg gradient *in vivo*, we predict that overscaling phenotype will be lost in *ds*>*vg*^*RNAi*^ discs. We sorted the discs into four groups according to their DV length (Fig. S13B) and quantified the patterns of Vg as we did for the wild-type discs (Fig. 3A-D). As predicted, on average the overscaling property of the Vg pattern is mostly lost (Fig. 3E, compare to Fig. 1G). Although there is a small, but positive slope when we plotted the relative width of the Vg pattern *vs*. DV length in these discs (Fig. 3F), this could be attributed to the fact that *vg*^*RNAi*^ expression does not result in a full knock-out of Vg-expression.

**Figure 3.**
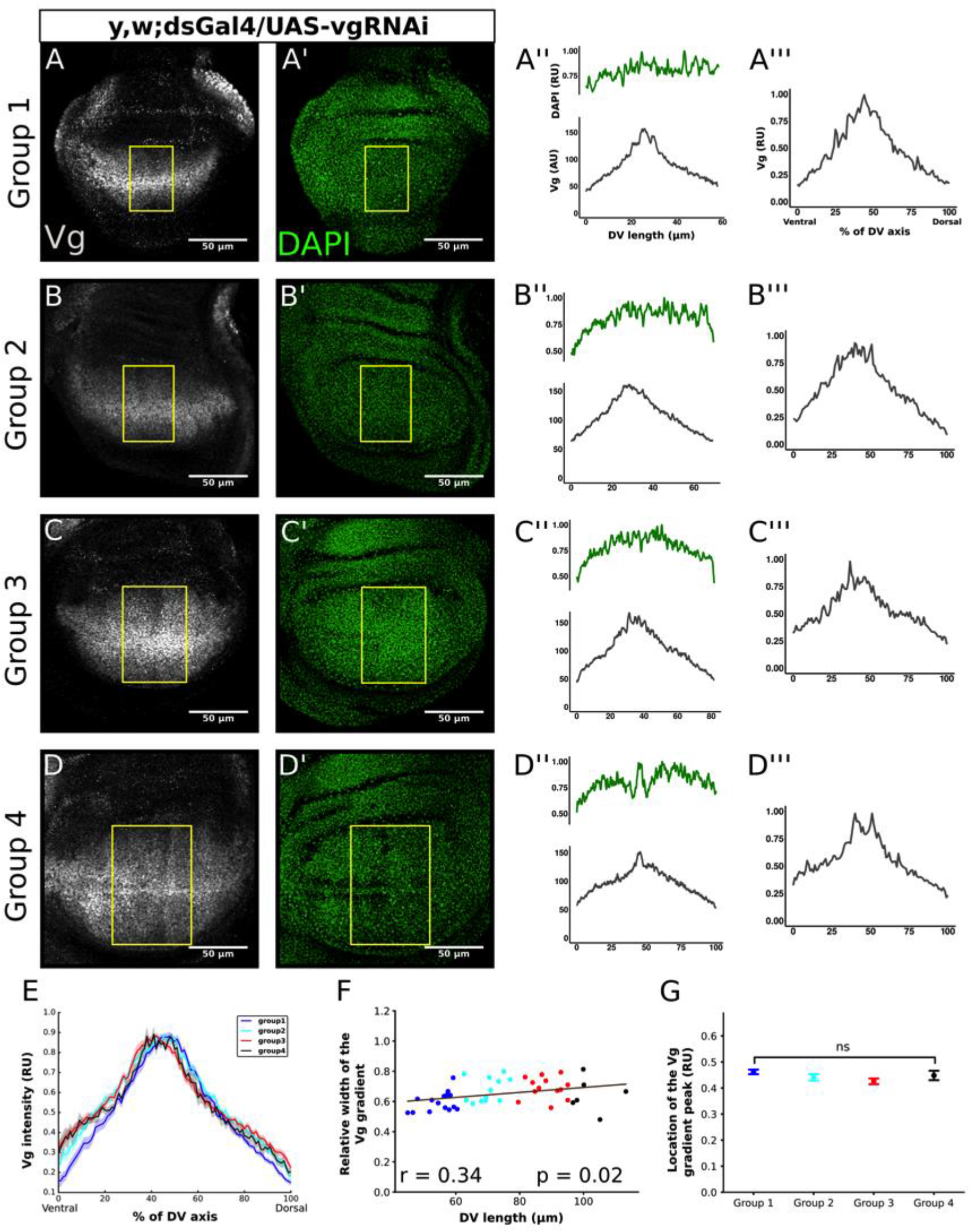
Genetic impairment of cell recruitment by *ds*>*vg*^*RNAi*^ results in almost perfect scaling of the Vg pattern. (A-D) Representative images of third-instar wing discs in which cell recruitment was blocked by expressing *vg*^*RNAi*^ in the domain of *ds* using the Gal4-UAS system. The discs were classified in four groups according to their DV length as defined in Fig. 1B. Discs immunostained with a Vg antibody (A-D) and DAPI (A’-D’) and the quantification of the Vg patterns in a rectangular region defined as in Fig. 1A in AU (A’’-D’’) and RU (A’’’-D’’’) units as in Fig. 1. The quantifications of the Vg patterns in A’’’-D’’’ are normalized to DAPI levels. (E) Median of the normalized Vg profiles (dark line) and SEM (shaded area) from all the discs in each group (n=15, 12, 13 and 6 for Groups 1, 2, 3, and 4, respectively). (F) Relative width of the Vg gradient (as defined in Fig. 1H) as a function of DV length; each dot in the graph corresponds to a different disc and is color-coded as in E. The solid line shows the linear regression of the data and the Pearson Correlation Coefficient, r, of the regression is displayed. The p-value corresponds to a Student-t statistical test assuming a zero-slope of the regression line as a null hypothesis. (G) Comparison of the location of the Vg gradient peak in RU for the discs in each group; error bars correspond to the SEM. (Statistical significance is analyzed using a Kruskal-Wallis non-parametric test; ns, p-value=0.075).

We considered if the loss of Vg overscaling in *ds*>*vg*^*RNAi*^ discs could result from an alternative mechanisms. For example, *ds*>*vg*^*RNAi*^ discs may experience higher cell proliferation or lower apoptosis rates at the edges of the pouch due to cell competition or changes in the slope of the gradient (Baéna-Lopez and García-Bellido, 2006). We also examined experimentally if the expansion of the Vg pattern is the result of oriented cell divisions. Since Vg inhibits *ds* expression transcriptionally (Zecca and Struhl 2007a) and the Fat-Ds pathway dictates the orientation of cell divisions along the proximal-distal direction (Baena-López *et al.* 2005), it is possible that the orientation of cell divisions could be affected in *ds*>*vg*^*RNAi*^ discs, resulting in a narrower Vg gradient. However, we found that cell proliferation, apoptosis, and the patterns of cell divisions are very similar in control and *ds*>*vg*^*RNAi*^ discs (Fig. S6 and S14). We conclude that preventing Vg expression in the *ds* expression domain blocks the recruitment-dependent overexpansion of the Vg pattern.

### A time-dependent cell recruitment process contributes to the Vg pattern during normal development

So far, the evidence of a cell recruitment mechanism that patterns the Vg gradient has been indirect. In order to investigate more directly the dynamics of the cell recruitment process during normal development, we used a recently-developed tool of dual-color fluorescent reporters, known as TransTimer (He *et al.* 2019), to examine the dynamics of the *vg*^*QE*^. The TransTimer expresses a rapid yet unstable GFP and a more stable, but slower RFP under *vg*^*QE*^Gal4 control. Hence, recently-recruited cells will express GFP but not RFP (or have a RFP to GFP ratio below a threshold), whereas cells that have been activating the reporters for longer than 12 hours will express RFP as well (Fig. 4A). We aimed to use this system, both to provide direct evidence of the recruitment process and to show that it takes place at a particular time during normal development, as revealed by our time course of Vg expression (Fig. 1G). Again, we used DV length as defined in Fig. 1 as a measure of developmental time and we quantified the reporters of the TransTimer under the control of *vg*^*QE*^Gal4. We found GFP-exclusive expression in discs that have a DV length in the 60-80 sm range (Fig. 4B-B’’’), whereas discs larger than 80 mm rarely exhibit GFP-exclusive pixels (Fig 4C-C’’’ and 4D-D’’’). This demonstrates the existence of cells that respond to the *vg*^*QE*^ and that this process temporally correlates with the major overexpansion of the Vg pattern, *i.e.* between groups 1 and 3 (see Fig. 1B,G). Altogether, our data directly reveal that during normal wing disc development, the Vg pattern is shaped by the recruitment of new cells at a specific period of time between early and mid third-larval instar.

**Figure 4.**
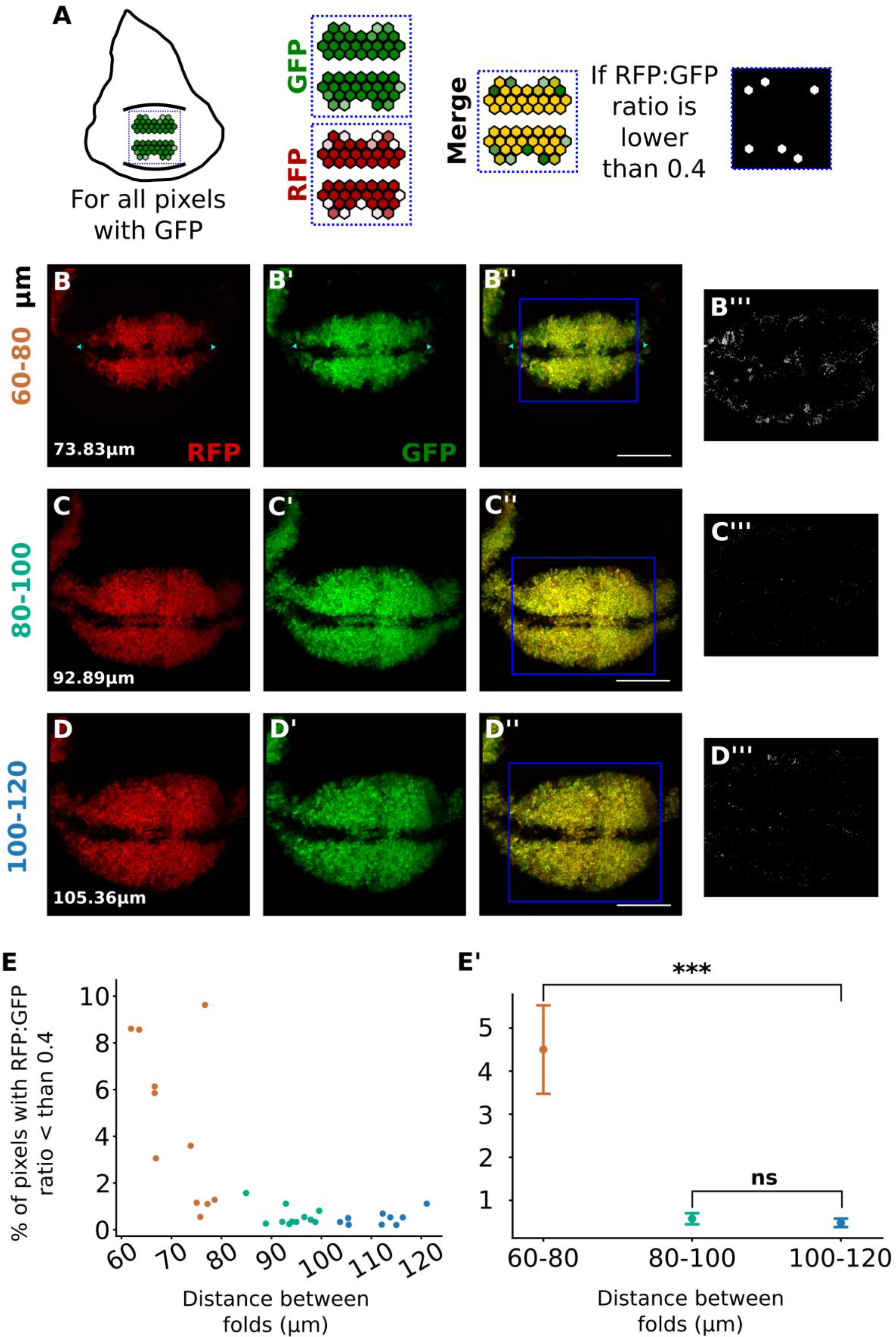
*vg*^*QE*^ is activated in new cells during early-to-mid third instar. Fixed wing discs from *vg*^*QE*^Gal4, UAS-TransTimer larvae (see Materials and Methods). (A) Scheme of the quantification approach. Only GFP-positive pixels within the wing pouch were considered in the analysis (see Materials and Methods). For each of these pixels, we looked at the corresponding value in the RFP channel and computed the RFP to GFP ratio; ratio values below 0.4 (see Materials and Methods) were taken as pixels coming from cells that recently activated the *vg*^*QE*^. (B-D) Representative discs of different DV length groups that correspond to different developing times (distance between folds is shown in white for each disc), that express RFP (B-D), GFP (B’-D’) or both (B’’-D’’). Some GFP positive cells in the most anterior and posterior locations are pointed with a cyan arrowhead in B-B’’. The pixels that have a RFP to GFP below the 0.4 threshold are depicted in white in B’’’-D’’’. (E, E’) A quantification of the percentage of the pixels with the RFP to GFP ratio below 0.4 in all the GFP positive pixels individual discs (E) and grouped by their DV length (E’). Kruskal-Wallis and Mann-Whitney U non parametric tests were used. ***, p-value = 0.00017; ns, p-value < 0.01.

### Impairment of cell recruitment results in smaller, but well-proportioned adult wings

We then asked whether cell recruitment could have an effect on adult wing size. Therefore, we compared *ds*>*vg*^*RNAi*^ vs. control adult wings (Fig. 5). Most adult wings of *ds*>*vg*^*RNAi*^ animals show the normal vein patterns (Fig. 5A-B). However, *ds*>*vg*^*RNAi*^ wings are on average smaller than control wings (Fig. 5C). We assume that cell recruitment takes place in dorsal- and ventral-most regions of the wing pouch, which correspond to proximal regions of the adult wing. Therefore, we predicted that *ds*>*vg*^*RNAi*^ adult wings would be smaller in proximal regions, but unaffected in distal regions. However, we found that representative areas of both proximal (Fig. 5D, inset) and distal (Fig. 5E, inset) regions of the adult wing are similarly reduced in *ds*>*vg*^*RNAi*^ animals (17 and 21 %, respectively; Fig. 5D-E). Furthermore, we found that *ds>vg^RNAi^* wings maintain their proximal-distal (longitudinal) to anterior-posterior (transversal) proportions, as the ratio of longitudinal to transversal dimensions is not statistically different between control and mutant adult wings (Fig. 5F). Taken together, we conclude that the impairment of cell recruitment in *ds*>*vg*^*RNAi*^ animals significantly affects adult wing size but not its proportions.

**Figure 5.**
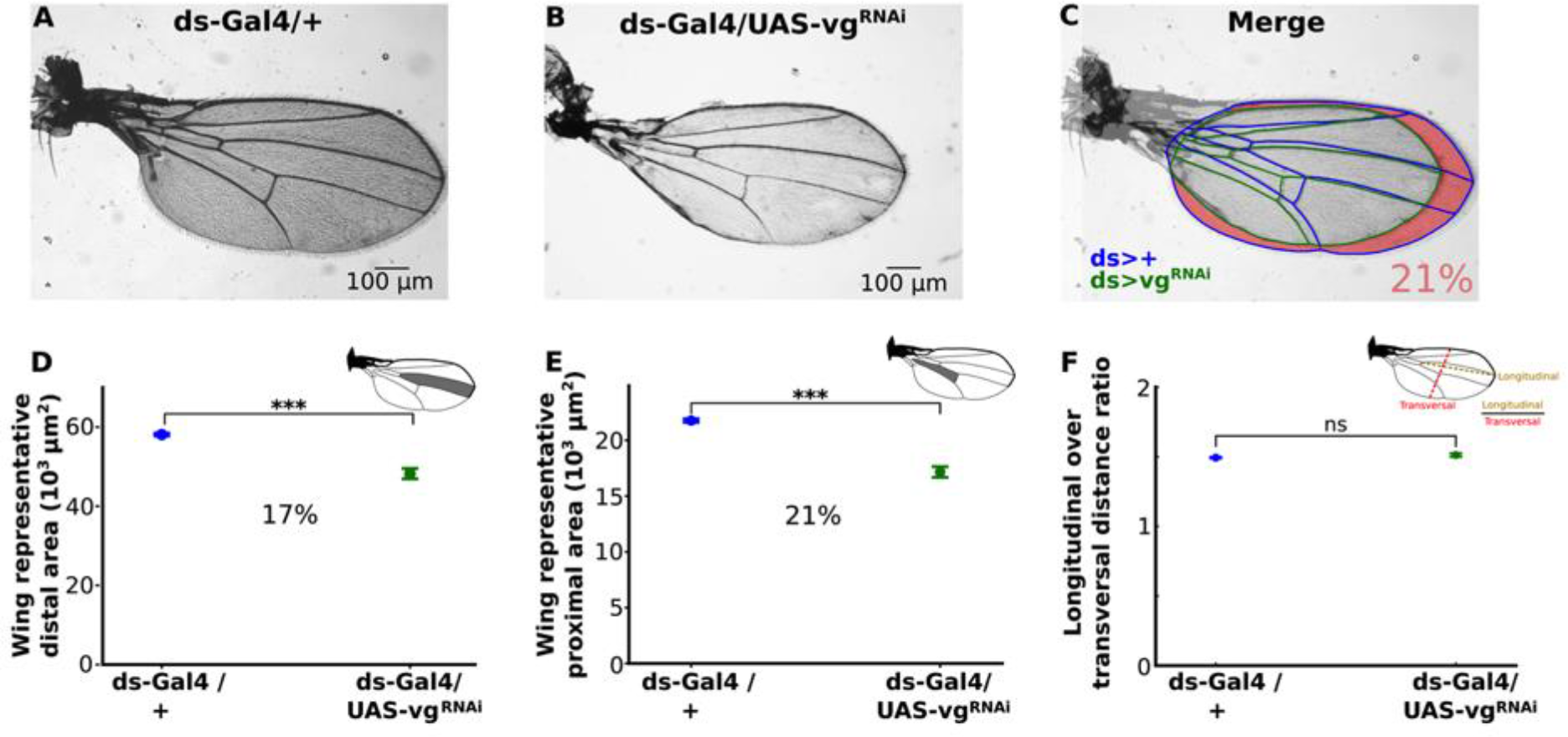
Genetic impairment of cell recruitment results in proportionally smaller adult wings. (A and B) Representative adult wings of *ds*>*vg*^*RNAi*^ (A) and dsGal4/+ (control, B) animals. (C) Merged wings shown in A (veins and margin colored in blue) and B (veins and margin colored in green). The area difference is shaded in pink. On average, control wings are 21% smaller than when recruitment is impaired. (D, E) Comparison of representative (see insets) distal (D) and proximal (E) areas in dsGal4/+ vs. *ds*>*vg*^*RNAi*^ wings. (Average percentage of reduction is shown, statistical significance is analyzed using a Mann-Whitney non-parametric test; ***, p-value<10-5). (F) Comparison of the ratio of longitudinal to transversal distance in dsGal4/+ vs. *ds*>*vg*^*RNAi*^ wings (Statistical significance is analyzed using a Mann-Whitney non-parametric test; ns, p-value<0.01).

## Discussion

How gene expression patterns that determine cell fate are established and contribute to organ size are important questions in developmental biology. There are two predominant models by which gene expression patterns are coordinated with organ growth. In the first model, patterns continuously depend on morphogen signaling; when the morphogen gradient changes, the patterns also change accordingly. In particular, if the morphogen scales to organ size, the patterns that depend on the morphogen would maintain their proportions relative to the final size. In the second model, a morphogen gradient establishes pre-patterns of gene expression that are then locked by positive feedback regulation. Once cells are locked in a determined fate, the final patterns are no longer dependent on the morphogen but are determined by organ growth. Particularly, when growth is uniform, this results in patterns that scale with size throughout development. In both of these models, the patterning process adjust to organ growth and the final patterns are invariant to organ size. Our data on patterning of the wing selector gene, *vg*, in the Drosophila wing disc support an additional conceptual model. After being established by Notch and Wg signaling, Vg expression is maintained through cell divisions by a mechanism involving Polycomb/Trithorax Responsive Elements. In a tissue that grows uniformly like the wing disc, this would result in a pattern that scales with tissue size. However, we find that the Vg gradient overscales relative to DV axis length (Fig. 1). This cannot be explained by non-uniform cell proliferation, apoptosis, or a substantial change in pattern of oriented cell divisions (Figs. S5, S6, S14). Instead, our modeling results suggest that Vg overscaling could be explained by a cell recruitment mechanism (Fig. 2). Experimentally, when cell recruitment is genetically blocked, the overscaling phenotype is mostly lost (Fig. 3). We provide direct evidence of the cell recruitment dynamics during Vg patterning (Fig. 4). Finally, we show that cell recruitment contributes to about 20 % of adult wing size (Fig. 5).

Based on qualitative evidence on Vg expansion in genetic mosaics, Zecca and Struhl identified the molecular mechanisms of recruitment and proposed that Vg-dependent cell recruitment could contribute to growth of the Drosophila wing disc (Zecca and Struhl, 2007a; Zecca and Struhl 2010). However, it remained unclear when and to what extent cell recruitment actually contributes to normal patterning and growth of the disc. Our study directly shows that cell recruitment works by overexpanding the endogenous pattern of a selector gene, thus contributing to both patterning and organ growth. To our knowledge, this is the first study where recruited cells are directly detected during wild-type development and where its contributions to pattern formation and organ growth are quantitatively determined.

Wg signaling is absolutely required for Vg expression and propagation along the DV axis (Zecca and Struhl, 2007a). But is Wg signaling the determining factor of the range of Vg expression? By overexpressing the Wg receptor, Frizzled2 in individual cells, a recent study has shown that the maximum range of Wg signaling is 11 cells from the DV boundary (Chaudhary *et al.* 2019). We determined the position of cells that acquire new expression of the GFP reporter in the TransTimer system and show that these cells are located 14-16 cells away from the DV boundary (Fig S15). How can we conciliate the requirement of Wg signaling for cell recruitment with the observation that cells are being recruited outside the reported range of Wg signaling? We think that *vg* expression may depend on memory of earlier, pouch-wide Wg signaling activity (Alexandre *et al.*, 2014). If this is the case, Wg signaling would only be a permissive factor and would not play a leading role in establishing the range or overscaling of Vg patterning.

In this study, we focused on Vg patterning along the DV axis, but other signaling pathways participate in Vg expression. For example, Decapentaplegic (Dpp), a member of the BMP family, patterns the disc along the anterior-posterior (AP) axis and has been proposed an input signaling for *vg* expression (Kim *et al.*, 1996). Importantly, our analysis using the TransTimer system does shows new activation of the GFP reporter in cells located in the anterior and posterior ends of the disc (arrowheads in Fig. 4B’’). Therefore, how Dpp signaling contributes to cell recruitment along the AP axis and participates in the 2-dimensional Vg pattern is a promising perspective left to future work.

Our TransTimer experiment shows that new cells are being incorporated into the *vg*^*QE*^ pattern, *i.e.* new cells get recruited, and reveal that this recruitment process is temporally controlled (Fig. 4). This is consistent with the major overscaling of the Vg width that occurs in discs of similar sizes in Fig. 1G. However, it is likely that the TransTimer reporters only show the earliest manifestation of the recruitment process and that this continues over the rest of the third larval instar by increasing the levels of Vg until they reach an appropriate level for wing-fate differentiation. Thus, we interpret the dynamics of Vg pattern formation as a two-step recruitment process: The first step takes place between early and mid third-instar, and results in the overscaling of the Vg gradient width (Fig. 1G,H). The fact that this step occurs during a narrow window of time (Fig. 4), suggests that the Ft-Ds polarization signal can propagate several cells without the need of a cell-by-cell expansion of Vg expression, *i.e.* several layers of cells could begin to be recruited at once (Wortman *et al.* 2017). The second step extends from mid to late third-larval instar and results in increasing levels of Vg at the tails of the gradient. This is captured by a decrease in the slopes of the Vg gradient in our time-course examination of wild-type Vg patterns (Fig. 1G and Fig. S4). The fact that the slopes at the tails of the Vg gradient never completely flatten suggests that the rates of Vg upregulation are position-dependent, possibly dictated by the strength of the Ft-Ds polarization which decays from its source near the DV boundary towards ventral- and dorsal-most positions. This two-step model of cell recruitment suggests that the spatial range of the process is limited by the length-scale of the Ft-Ds polarization signal or by the availability of recruitable cells within the wing pouch.

We show that cell recruitment contributes to approximately one fifth of the total adult wing size (Fig. 5C). An interesting question that remains is to understand the purpose of cell recruitment as a growth mechanism. In other words, what is the advantage of this developmental design over simply adjusting cell sizes or cell proliferation rates to achieve a target final size? It is possible that cell recruitment plays a role in conferring some sort of robustness to developmental growth control. For example, perhaps recruitment compensates for variations in cell proliferation and could explain why final wing disc size is robust to perturbations against cell proliferation rates or cell size (Day and Lawrence, 2000).

Recent studies have showed that a cell recruitment mechanism is present in early thyroid and kidney development in mammals (Fagman et al. 2006; Lindström et al. 2018), but it is unclear how much recruitment contributes to organ growth or what are the molecular players of the recruitment signal in these contexts. Given the widespread conservation of the recruitment signal components that operate in the Drosophila wing, it would be interesting to explore whether the homologous cell recruitment signal operates as a growth-by patterning mechanism in other developing organs.

## Materials and Methods

### Fly stocks and genetics

The following stocks of *Drosophila melanogaster* were used: *y*, *w* provided by Fanis Missirlis (Cinvestav, Mexico); UAS-TransTimer provided by Li He (Norbert Perrimon’s Lab, Harvard Medical School, USA); dppGal4 (Bloomington Drosophila Stock Center, BDSC #1553); Act>STOP>LacZ (BDSC #6355); UAS-hid (BDSC #65403); UAS-*vg*^RNAi^ (Vienna Drosophila Resource Center # 16896); ds-Gal4, UAS-GFP (II) provided by Gary Struhl (Columbia University, New York, USA); vg^QE^-Gal4 (III) (BDSC #8229). Imaginal discs were dissected from third-instar larvae of both sexes. For normal endogenous Vg patterns quantification, the *y, w* stock was used as wild-type flies (Fig. 1) and kept at 25°C during egg laid and development. For recruitment-impairment experiments (Figs. 3 and 5), dsGal4, UAS-GFP flies were crossed to UAS-*vg*^*RNAi*^ (for recruitment inhibition) or the *y, w* stock (control) and were kept at 25°C during egg laid and then changed to 29°C during development to increase Gal4 system efficiency. For the evaluation of the *vg*^*QE*^ dynamics (Fig. 4), UAS-TransTimer reporter, a destabilized GFP (dGFP) protein combined with a stable RFP (He *et al.* 2019), was placed under vg^QE^Gal4 driver to quantify cells that recently activated this enhancer. Taking advantage of the different maturation rates of both proteins (dGFP about 0.1 h, and RFP about 1.5 h), we selected all the GFP pixels with intensity values above a threshold in the wing pouch for each disc; this threshold was set by ranking all the pixels within by their GFP intensity in three different groups using the machine learning K-Means function from scikit-learn python package (https://scikit-learn.org/stable/index.html) and all the pixels in the lower group were taken away. Next, we looked for the corresponding RFP channel and computed a ratio of RFP over GFP intensity. If this ratio was lower than 0.4 at a specific pixel (this value corresponds to the half maximum for GFP taken from Fig. 3b in He *et al.* 2019), we marked this pixel and considered it as new activation of the *vg*^*QE*^ (Fig. 4A).

### Immunostaining

After dissection in a stereoscopic microscope (Nikon SMZ800), discs were fixed in PEM-T (PEM with 0.1% of Triton X-100) with 4% paraformaldehyde, washed 3 times and blocked in PEM-T with 0.5% of BSA (Bovine Serum Albumin) for 2 hours at room temperature. Then, samples were stained with primary antibodies at 4°C overnight at the following dilutions: rabbit anti-Vg (a gift from Sean Carroll and Kristen Guss, 1:200) and rabbit anti-Caspase 3 (Cell Signaling Technology, Cat# 9661, 1:200), mouse anti-beta Galactosidase (Promega, Cat #Z378A. 1:1000). DAPI (1:1000) was used to stain nuclei. 5-ethynyl-2’-deoxyuridine (EdU) labeling was performed using the Click-iT EdU Alexa Fluor 647 Imaging Kit (Invitrogen Cat#C10340) following manufacturer instructions. Primary antibodies were detected with Alexa Fluor 488 anti-mouse and 647 anti-rabbit (1:1000). Imaging was done with a confocal microscope (Leica TCS SP8 Confocal Microscope) using a 63X oil-immersion objective.

### Wing mounting

Adult flies were dehydrated overnight in 70% ethanol and then separated by gender. Wings were dissected in 50% ethanol. The isolated wings were mounted and dried in a plate at 60 °C. Imaging of adult wings was done using a bright-field microscope (Nikon eclipse Ci-L/Ci-S) using a 4X objective. Wings were dissected from female and male flies and were independently analyzed.

### Quantification and Statistical Analysis

The details about the quantification of Vg (Figs. 1 and 4) are provided in the Supplementary Material. All the quantifications were performed using Python 3 programming language (https://www.python.org/download/releases/3.0/). The Python packages used for quantification and statistics were NumPy (http://www.numpy.org/), pandas (https://pandas.pydata.org/), and SciPy (https://www.scipy.org/). The statistical methods used were T-test or Mann-Whitney test for comparing pairs of parametric or non-parametric datasets, respectively, and Kruskal-Wallis for more than two non-parametric datasets.

### Wing disc and adult wing quantification

Vg patterns in the wing disc were quantified as explained in Fig. S1. Wing representative distances and areas were quantified using the straight line and polygon selection tools respectively in ImageJ/Fiji software (https://imagej.net/).

### Simulations

All simulations were performed using Morpheus 1.9.3 software (https://imc.zih.tu-dresden.de/wiki/morpheus). The time dependence of the Vg concentration was modeled by means of an ordinary differential equation for each cell (Fig. 2A), and the tissue was modeled by a Cellular Pots Model (CPM). Examples of the .xml files used to perform simulations are provided as supplementary files.

### Image processing and data visualization

All the images were processed and analyzed using ImageJ/Fiji software, the matplotlib (https://matplotlib.org/), pandas (https://pandas.pydata.org/), and NumPy (http://www.numpy.org/) python packages. The data, after being analyzed, were visualized with the seaborn (https://seaborn.pydata.org/) and Matplotlib python packages.

## Acknowledgments

We thank Li He from Norbert Perrimon’s Lab for kindly providing us the UAS-TransTimer flies and Fanis Missirlis for critical comments on the manuscript. We also thank the members of the Nahmad and Chara laboratories for interesting discussions.

