## Supplementary figures and images for "A dynamic cell recruitment process drives growth of the Drosophila wing by overscaling the Vestigial expression pattern"

### Video S1

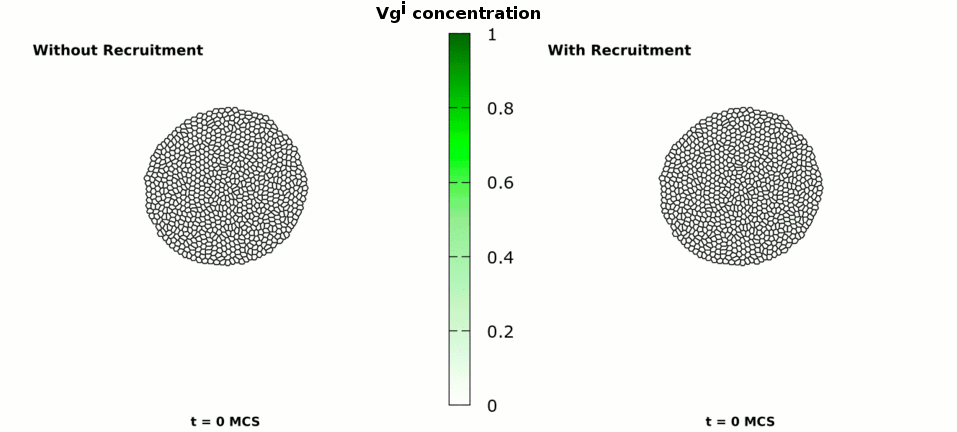
