## Supplemental Information for "A dynamic cell recruitment process drives growth of the Drosophila wing by overscaling the Vestigial expression pattern"

### 1 - Spatio-temporal quantification of Vg pattern in wing discs

The Vg pattern in each disc was obtained by quantifying the Vg intensity of a maximum projection image of the z-stack confocal images. We then used the line selection tool in ImageJ (ImageJ RRID: SCR_003070) from the ventral to dorsal fold (yellow rectangle in Fig. S1A). The width of the line (in pixels) was taken to be 0.6 of the distance between the folds (converted to pixels) so that we can get an average pattern along the DV axis (yellow rectangle in Fig. S1A).

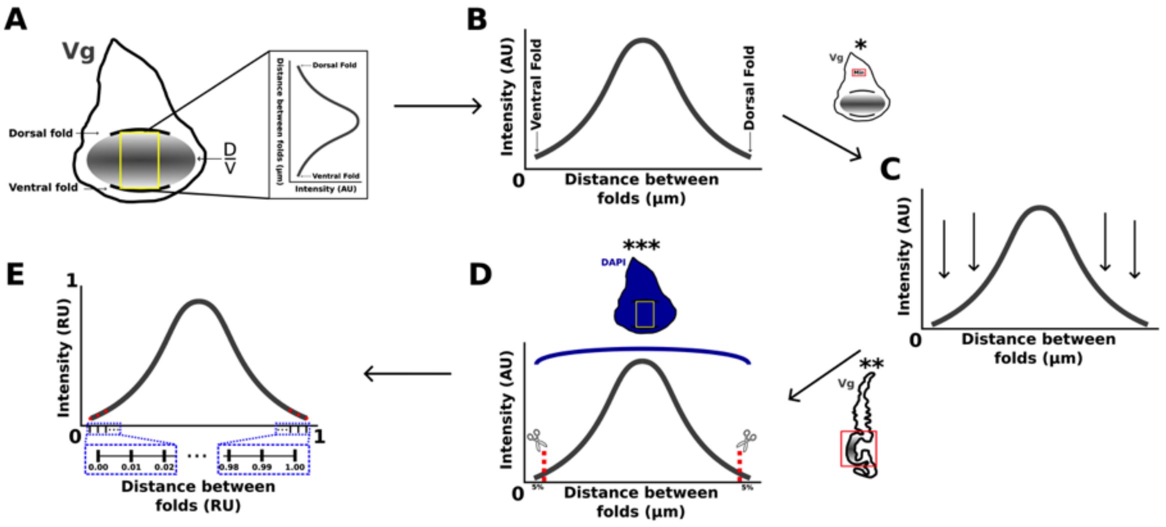


**Figure S1. Outline of Vg quantification in wing discs.** Raw data from confocal images stained for Vg and DAPI were processed as follows: (A-B) Three-dimensional data were converted into a Vg profile first by obtaining a maximum projection and then collecting the average intensity along vertical lines of pixels within an area of interest (yellow rectangle in A). This gives a Vg profile in arbitrary units (AU) as a function of DV position from the ventral to the dorsal fold (in µm, B). (C) Background levels of Vg from a non-pouch area (*) were substracted to all positions in the Vg pattern. (D) Since the pouch area of the disc is not flat (** shows a cross section of the disc), we used DAPI (which we assumed is homogeneous, ***) as a normalizing signal (see Eq. S1-S4 in Supplemental Information. To avoid a high noise-to-signal ratio very close to the folds, we did not take into account the first and last 5% of data points of the Vg profile. (E) Background-corrected and DAPI-normalized Vg profile is plotted in relative units (RU) by dividing by the maximum Vg intensity and rescaling the distance between the folds into % of DV length (0%, ventral; 100%, dorsal).

Using the Plot Profile function in ImageJ we plotted the average intensity values of Vg within the selected line or rectangle against the distance between folds (in μm) obtaining a pattern of Vg intensity as a function of distance from the ventral to the dorsal fold ((Fig. S1B). We exported the data of the profile to continue with the analysis.

We imported the data in Python 3.6 programming language (Python, RRID:SCR_008394) and subtracted background levels of Vg staining to every Vg intensity value (Fig. S1C). To get a value of Vg background, we averaged Vg levels in a region of 50 x 50 μm located in the notum region of the disc where Vg is not expected to be expressed (* in Fig. S1).

Because the wing blade is not flat and is concave (** in Fig. S1), we wondered how much of the graded pattern of Vg is due to confocal signal attenuation. To address this technical issue, we made a correction with the nuclear marker DAPI, which we assumed is expressed uniformly in all nuclei of the disc (*** in Fig. S1). Let DAPI(real) be the true average expression of DAPI in every nuclei and α(Z) be the attenuation function of DAPI that is equal to 1 for Z=1 and decreases as a |function of the focal plane Z. Then, the observed values of DAPI at the Z plane are given by:

$$\mathrm{DAPI(Z)} = \mathrm{DAPI(real)}\ \alpha(\mathrm{Z}) \qquad \text{(Eqn. S1)}$$

After reordering the equation, we find that α(Z) is the normalized value of DAPI in every Z frame.

$$\alpha(\mathrm{Z}) = \frac{\mathrm{DAPI(Z)}}{\mathrm{DAPI(real)}} = \mathrm{DAPI}_{norm}(\mathrm{Z}) \qquad \text{(Eqn. S2)}$$

The real values of Vg should be given by:

$$\mathrm{Vg(obs)} = \alpha(\mathrm{Z}) \cdot \mathrm{Vg(real)} \qquad \text{(Eqn. S3)}$$

Substituting α(Z) with $DAPI_{norm}(Z)$ and reordering the equation we get that the real value of Vg is the observed value divided by the DAPI normalized value in that Z position that goes from numeric 0 to 1 values.

$$Vg(real) = \frac{Vg(obs)}{DAPI_{norm}(Z)} \qquad \text{(Eqn. S4)}$$

By doing this correction, we artificially "flatten" the wing blade. Because Vg became almost undetectable as the folds went deeper, the correction produced abnormal high levels of Vg at the folds, so we decided to subtract the final five percent of the data at each (dorsal and ventral) end (Fig. S1D red dotted lines).

Finally, in order to compare the Vg pattern curve between different discs, we normalized the Vg intensity and the distance between folds and divided the normalized distance in bins of 0.01 (Fig. S1E blue dotted rectangles), making an average of all the Vg values per bin and resulting in 100 values of Vg between 0 and 1 for each disc (Fig. S1E).

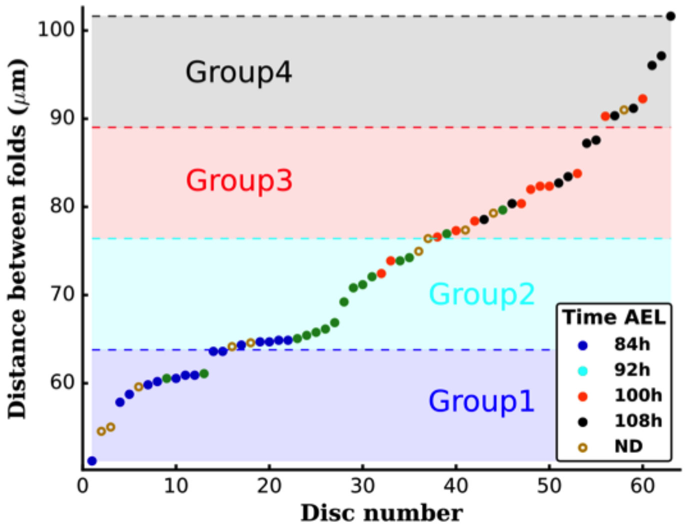


**Figure S2. DV length is correlated with disc age.** Embryos from 0-6 h were collected and larvae were fixed at 84, 92, 100 and 108 h AEL at 25 ˚C. In this figure, which is equivalent to Fig. 1B, each disc is color-coded by their age. (ND corresponds to a sample that were not aged, but included in the analysis of Fig. 1). Note that there is a clear correlation between groups (defined by distance between folds) and time AEL.

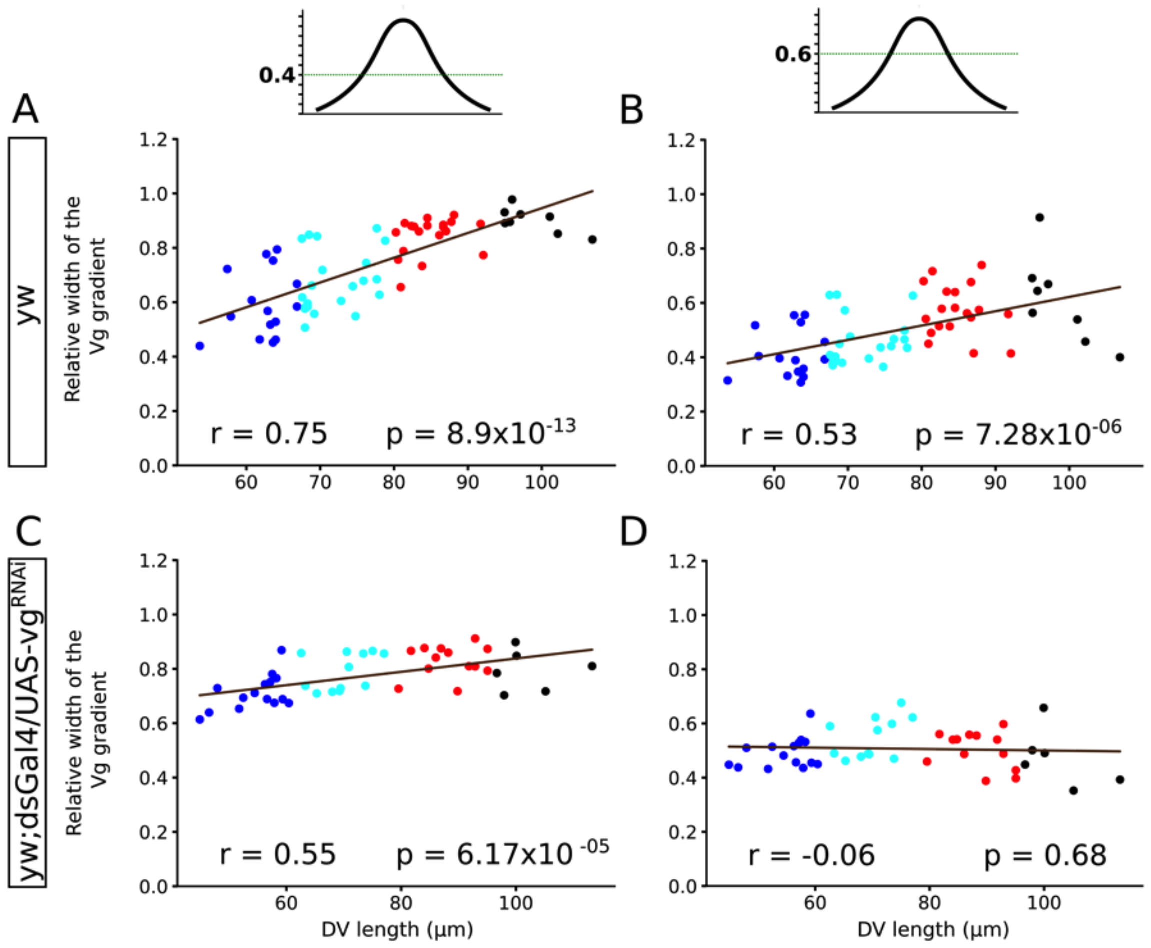


**Figure S3. Scaling properties of the Vg pattern is similar when using other thresholds to define the width of the pattern.** In the main text (Figs. 1 and 3) we used 0.5 to compute the widths of the Vg patterns (Fig. 1H and Fig. 3F). Here we show the relative width of the Vg (A, B in wild-type discs; C, D in recruitment-impaired discs) patterns when the width is defined at 0.4 RU (A and C) or 0.6 RU (B and D) as a function of DV length. The Pearson Correlation Coefficient (r) and a p-value are shown for each plot. Each dot corresponds to a single disc and discs are color-coded according to the group they belong as in Fig.1G (Vg in wild-type) and Fig. 3E (Vg in recruitment-impaired wing discs).

### 2. Investigation of the Vg pattern by analysis of slopes

Since the shapes of the Vg pattern do not fit a standard mathematical function (such as a Gaussian), we opted for investigating how the slopes of the pattern change at different locations (Fig. S3). We applied a linear regression model, using the polyfit function of the numpy module (NumPy, RRID:SCR_008633) in python 3.6, for different intervals or regions in all the normalized Vg patterns, and captured the resultant slope to compare the Vg patterns among groups.

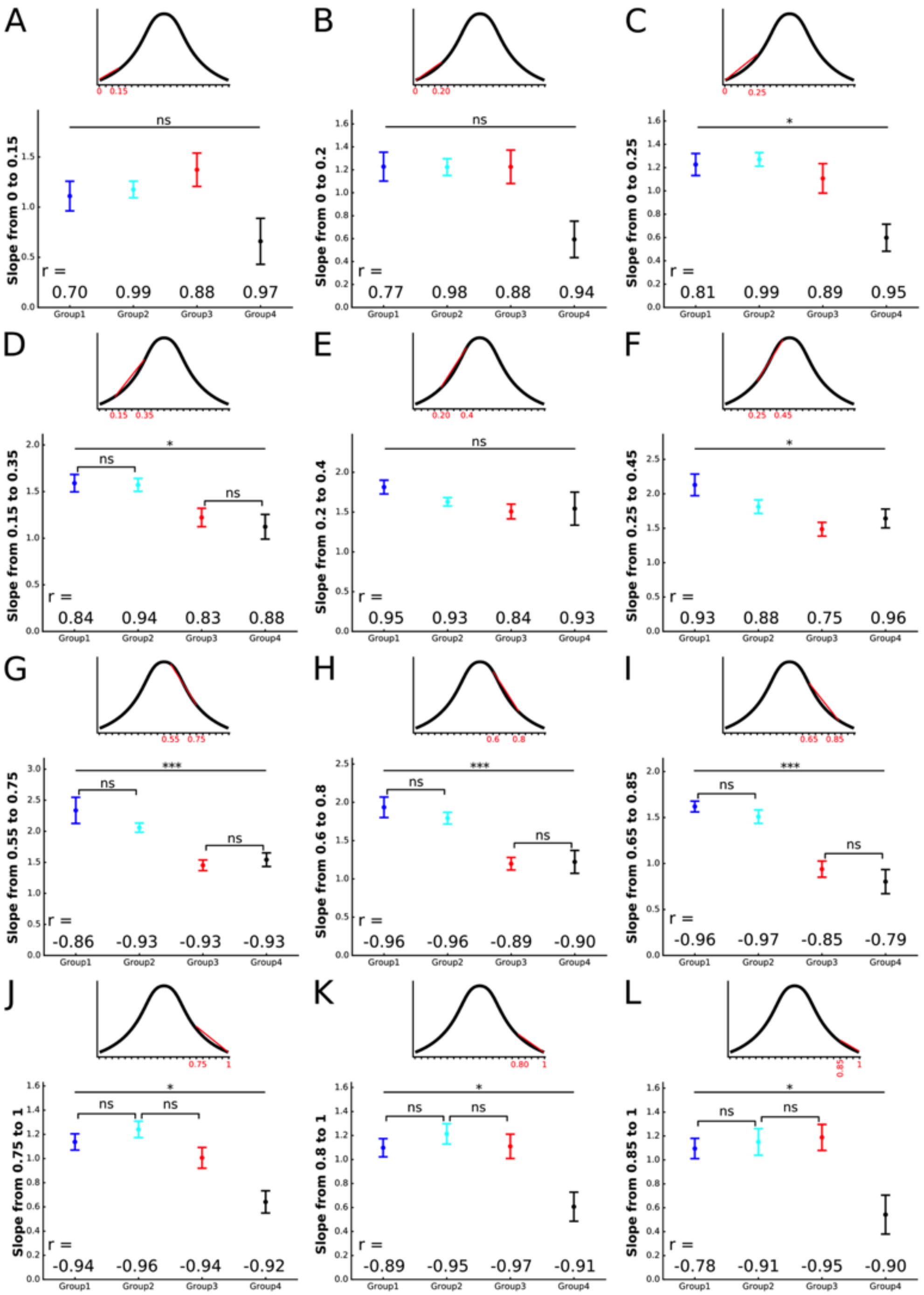

**Figure S4. Linear regression analysis for slope characterization of the Vg patterns in wild-type discs.** Comparison of slopes of the linear regression model calculated from the wild-type Vg patterns in relative units at different locations (as shown in the insets above each panel) among groups. Note that the results are independent of the choice of the specific interval. Statistical analysis was performed using the Kruskal-Wallis non-parametic test. Pairwise statistical comparisons between groups were done using a Mann-Whitney non-parametric test (ns, p-value>0.01; *, p-value<0.01; ***; p-value<$10^{-4}$). The Pearson correlation coefficient value (r) for each linear regression analysis is shown.

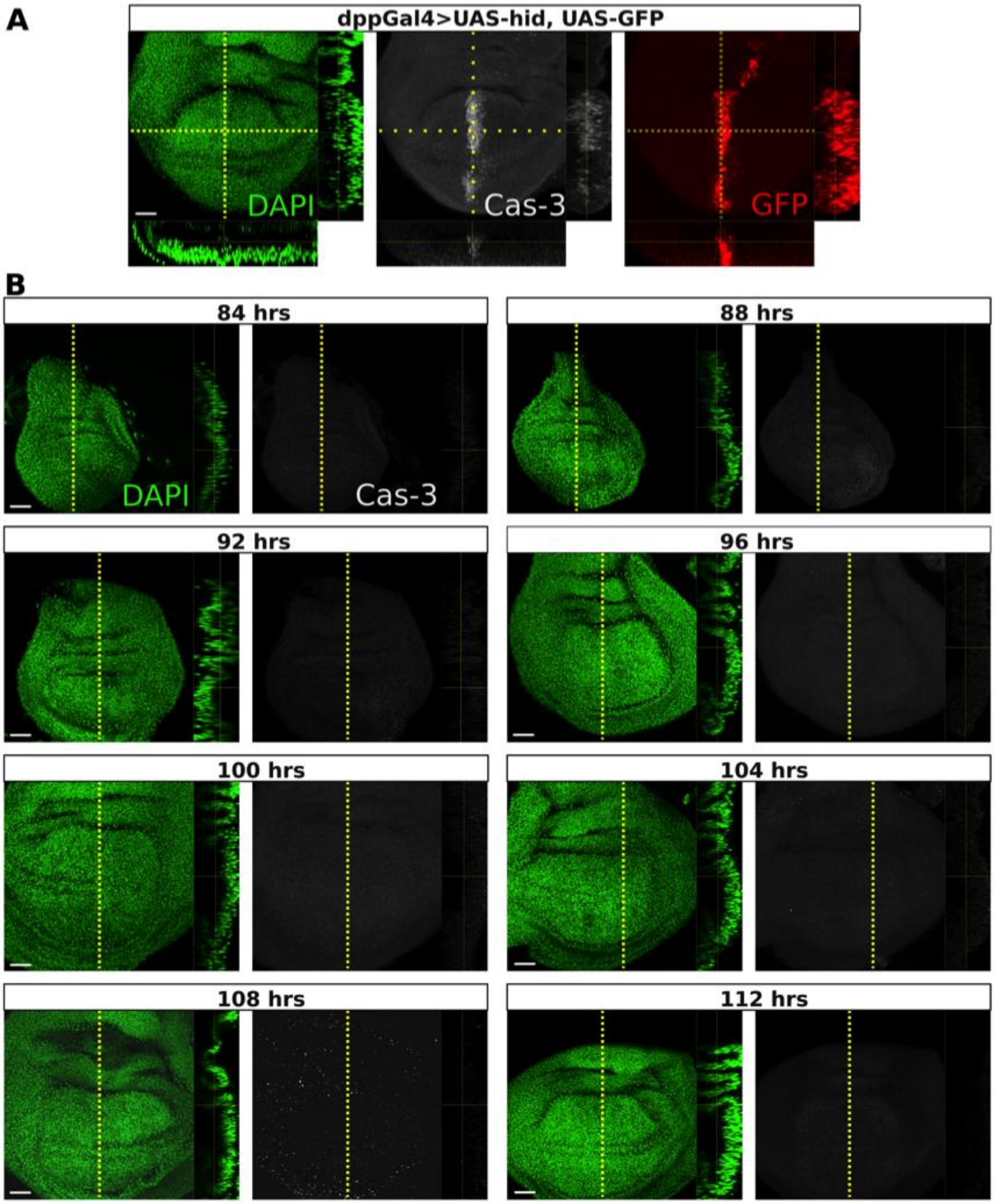


**Figure S5. Apoptosis occurs at low frequency and is nearly homogenous during wild-type development** (A) Discs immunostained with Caspase 3 (Cas-3) antibody (gray), DAPI (green), and GFP (red). The yellow dashed lines are drawn approximately halfway between the center and edge of the pouch, indicate sections along the AP boundary (right panel) oriented with dorsal up and apical right, and along DV border (bottom panel) oriented with apical down (A). (B) positive control disc undergoing ectopic induction of apoptosis by overexpressing the pro-apoptotic gene *hid* along the AP border (*dpp*-Gal4, UAS-GFP; UAS-*hid*). The following rows depict representative wild type wing discs at 84 h, 88 h, 92 h, 96 h, 100 h, 104 h, 108 h and 112 h AEL. Images in A and B were captured under the same imaging conditions. Scale bar: 10 μm.

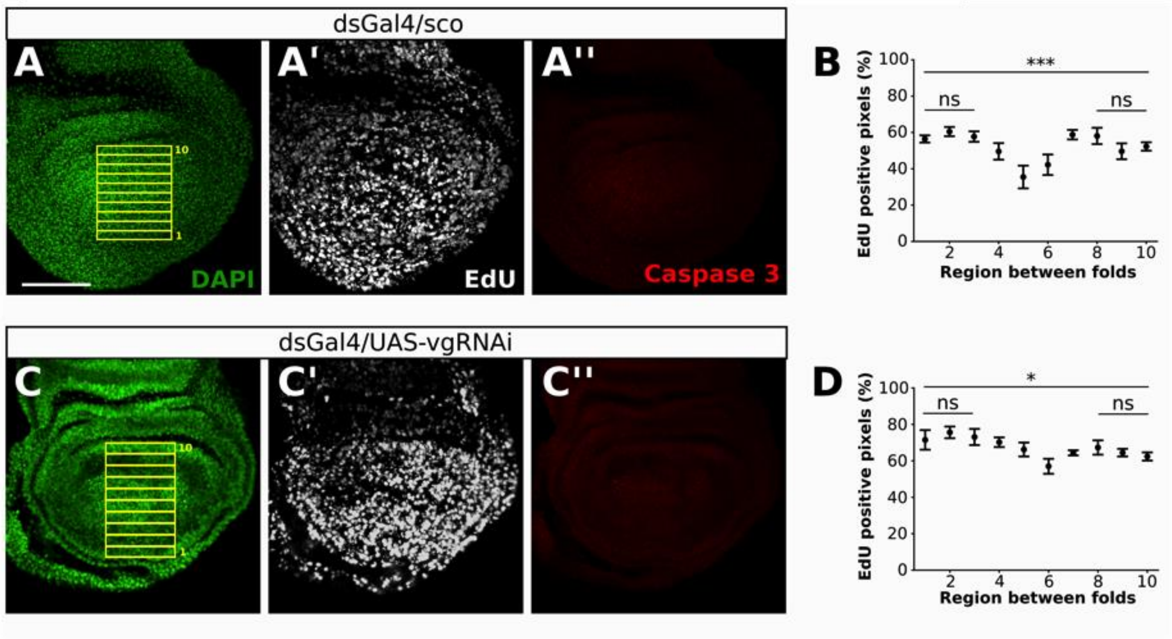


**Figure S6. Cell proliferation and apoptosis patterns are similar in *ds>vg$^{RNAi}$* and control discs.**
(A, C) Representative dsGal4/sco (control; A) and *ds>vg$^{RNAi}$* (C) wing discs in which DAPI is shown in green (A, C), cell proliferation is marked by EdU incorporation (gray; A', C') and Caspase 3 immunostaining is shown in red (A'', C''). (B, D) Quantification of cell proliferation in ten regions (yellow rectangles) from the ventral (1) to the dorsal (10) fold in a control (B) and *ds>vg$^{RNAi}$* (D) discs. Statistical tests were performed in all regions as well as in dorsal-most and ventral-most regions using a one-way ANOVA test; (*) corresponds to p-value<0.05, (***), p-value< $5x10^{-4}$, and non-significant (ns): p-value>0.05. n=7 (control), n=8 (*ds>vg$^{RNAi}$*). Note that significant differences in cell proliferation only occur around the DV border, but no differences were observed elsewhere. Scale bar: 50 µm.

### 3. Multiscale mathematical model of Vg distribution in the wing pouch

We modeled the distribution of Vg in the dorsal compartment of the wing pouch by means of a multiscale approach combining a Cellular Potts Model (CPM; Graner and Glazier, 1992; Glazier and Graner, 1993) (Fig. S7A-B), describing the cellular dynamics, with the following ordinary differential equation, accounting for the rate of change of Vg concentration in each cell *i* ($Vg^i$):

$$\frac{dVg^i}{dt} = \alpha M^i + \beta \frac{(Vg^i - Vg^i_{neighbor})^2}{\gamma^2 + (Vg^i - Vg^i_{neighbor})^2} - \delta Vg^i \qquad \text{(Eqn. S5)}$$

The first production term on the right hand side of the differential equation resembles the presence of a morphogen with a non-homogenous normalized profile $M^i$. This dependence of a morphogen accounts for the spatial-dependent activation of Vg by signaling factors in cells abutting the DV boundary including Notch and Wg. We will assume that the shape of this morphogen is given by the following non-scaling function:

$$M^i(y) = Ae^{-k\sqrt{(y-y_0)^2}} + B \qquad \text{(Eqn. S6)}$$

where $y- y_0$ represents the distance to the DV boundary (in μm). The parameters $A$, $B$ and $k$ were fitted to the average of normalized Wg profiles (Data not shown; n=32). The values of this fitting exercise are reported in Table S1:

| Parameter | Value (A.U.) |
|---|---|
| $A$ | 0.77 |
| $B$ | 0.23 |
| $K$ | 0.018 |

**Table S1.** Parameters values used in the simulations of Eqn. S5. The values are those that best fit the experimental profile of Wg to a curve of the form of Eqn. S6.

The second term of Eqn. S5 encodes our proposed recruitment mechanism (Fig. S7C), and it is given by a production term that depends on the difference in Vg concentration between the actual cell and the average of the concentration of its neighbors ($Vg^i_{neighbor}$) by a second order Hill function. Hence, this production term becomes relevant when the concentration of the cell is different from the one of its neighbors and negligible when they are similar (Fig. 2A). The third term in Eqn. S5 corresponds to a degradation process in a linear regime. When we considered a model without recruitment, the second term was not included in Eqn. S5.

We restricted our model to account for the Vg expression profile within the dorsal compartment of the wing pouch, because there are compartment-specific differences in the growth of the dorsal and ventral compartments (Fig. 1I). Taking into account these difference may require considering spatial differences between compartments, which we decided not to take into account. Also, the cells at the DV boundary were not taken into account in the comparison between experimental and simulated data because the model does not include additional effects exerted by the $vg^{BE}$.

The Cellular Potts Model (CPM) in 2D consists in a grid of *N* sites (Fig. S7A), in which each cell consists of *n* grid sites (*n* is equal to the cell surface), and has a membrane that consists in $n_{membrane}$ sites ($n_{membrane}$ is equal to cell membrane length/perimeter). In addition, cells are in contact with a surrounding medium, which has $n_{medium}$ grid sites. A Hamiltonian characterizing each configuration of the cell lattice depends on the adhesion between cells or with the medium, and on the elastic behavior of the cell area as well as its perimeter, as it can be seen in Eqn. S7:

$$H = \sum_{<i,j>} J_{ij}\left(1 - \delta_{ij}\right) + \sum_{i} \lambda_S (S(i) - S_T)^2 + \sum_{i} \lambda_P (P(i) - P_T)^2 \qquad \text{(Eqn. S7)}$$

where the first sum is over all pair of contacts between cells $i$ and $j$, and between cells and medium, $J_{ij}$ represents an energy per contact length due to adhesion between all sites in contact of cells $i$ and $j$. Second and third sums are over all cells, $\delta_{ij}$ is a Kronecker delta, $\lambda_S$ is an elastic surface constant and $\lambda_P$ is an elastic perimeter constant, both tending to lead cell surface $S(i)$ and perimeter $P(i)$ to their target values $S_T$ and $P_T$.

The system evolves following a Monte Carlo algorithm (Metropolis *et al.*, 1953). In each simulation step, for a grid of *N* nodes, *N* random attempts of elementary copy of a site to one of its neighbors are done. If this attempt minimizes the Hamiltonian, it is accepted. If not, it is accepted with a Metropolis acceptance probability $p = e^{\frac{-\Delta H}{T}}$, where $\Delta H$ is the energy gained with the copy attempt, and $T$ resembles a temperature (Fig. S7B). The initial condition of the model is such that Vg concentration of each cell is zero. All simulations were performed using Morpheus (Morpheus, RRID:SCR_014975) package (Starruß *et al.*, 2014).

### 4. CPM parametrization

The initial configuration consists in a quasi-circular distribution of 815 cells (Fig. S7A), centered in a lattice of 1250 x 1250 sites. The initial number of cells was determined fitting our model to the experimental data (see next section).

We scaled the neighborhood used to estimate the boundary length of CPM shapes by size, with a 3rd order neighborhood (to promote hexagonal cell shapes), and sampled the lattice sites for elementary copy with a uniform random distribution over all lattice sites.

We selected the parameters of the CPM Hamiltonian as follows (Table S2):

| Parameter | Value |
|---|---|
| Temperature (T) | 1.0 A.U. |
| Cell-cell interaction ($J_{cell-cell}$) | 1.0 A.U. |
| Cell-medium interaction ($J_{cell-medium}$) | 1.0x10$^{-6}$ A.U. |
| Surface elastic constant ($\lambda_S$) | 110 A.U. |
| Perimeter elastic constant ($\lambda_P$) | 100 A.U. |
| Target surface ($S_T$) | 250 A.U. |
| Target perimeter ($P_T$) | 0.059 A.U. |
| Simulation time | 240 MCS |

**Table S2**. Parameter values of the CPM Hamiltonian

We selected interaction parameters in order to obtain a confluent rounded tissue, while we choose the relation between target surface and perimeter of the cells to obtain mostly hexagonal cell shapes. We choose the simulation time to get realistic tissue configurations and yet reasonable computational times. To simplify the model, we set the Temperature in 1 which induced a minimal rugosity of the cells within the simulation times adopted.

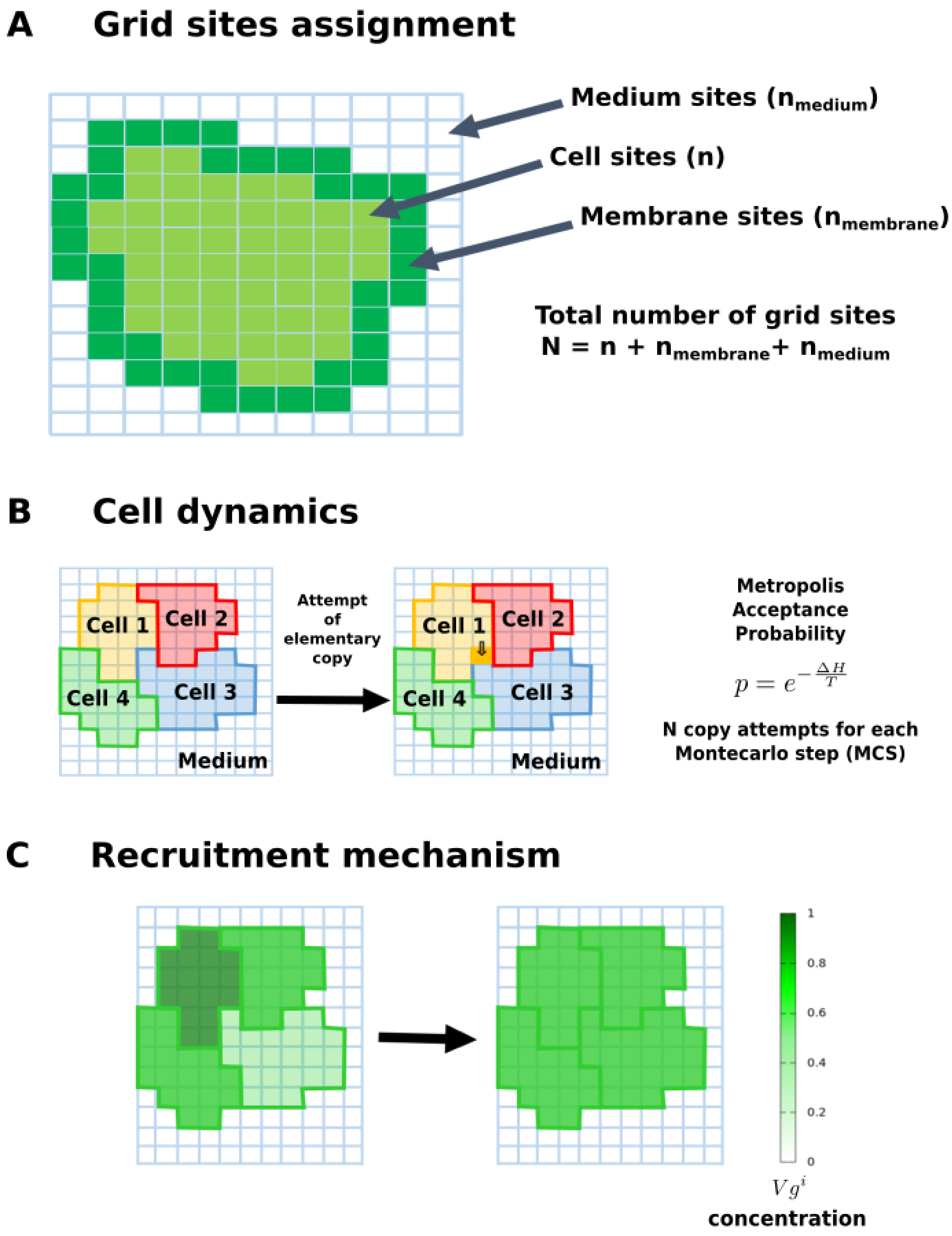


**Figure S7. Cellular level of the multi-scale model.** At the cellular level, we modeled the system by means of a Cellular Potts Model (CPM), in which each cell has a Vg concentration and senses the concentration from its neighbors. (A) The CPM consists of a lattice of *N* sites that can be part of cells or medium. (B) We modeled the evolution of the system, following a Metropolis-Montecarlo algorithm. (C) We quantitatively implemented the recruitment mechanism originally proposed in qualitative terms by Zecca and Struhl (Zecca and Struhl, 2007a). The Vg level of each cell depends on the Vg expression of its neighbors.

### 5. Model implementation of cell proliferation and its parametrization

We assumed that, in each simulation step, cells divide upon reaching the end of their cell cycle. Then, a segment along the cell minor axis is drawn, dividing the cell in two parts. The upcoming daughter cells evolve as the other cells in the tissue inheriting all the properties of the mother cell, except its perimeter and surface (Fig. S8A). To determine the cell cycle parameters together with the initial number of cells we followed the following procedure. We modeled a tissue in an initial circular configuration of $N_0$ cells and we assigned a position along the cell cycle to each cell according to a random uniform distribution within the interval [0, $\mu$] MCS. We then assumed that every daughter cell divides after a cell cycle length normally distributed with parameters $\mu$ and $\sigma$. Thus, we performed simulations varying the parameters $N_0$, $\mu$ and $\sigma$. We found that the values of the parameters $N_0$ = 815, $\mu$ = 127 MCS and $\sigma$ = 20 MCS allow the simulated tissue to reproduce the time evolution of the number of cells in the wing pouch (Fig. S8B) for the total simulation time (assuming that the four experimental groups were evenly distributed in time).

### 6. Fitting the models to the experimental data of Vg expression

In this article, two multi-scale models (with and without recruitment) were fitted to the here reported experimental data on the time course of the Vg profiles within the wing pouch (Fig. 1). We performed the fitting procedure varying the models' parameters values while minimizing a score function of the residuals between conveniently normalized experimental and simulated Vg profile data (see section 8 for more details). The model involving recruitment was explored in a 4-dimensional parameter space (given by the parameters $\alpha$, $\beta$, $\gamma$ and $\delta$, from Eqn. S5) while the model lacking recruitment was investigated in a 2-dimensional parameter space (consisting of only the parameters $\alpha$ and $\delta$, from Eqn. S5).

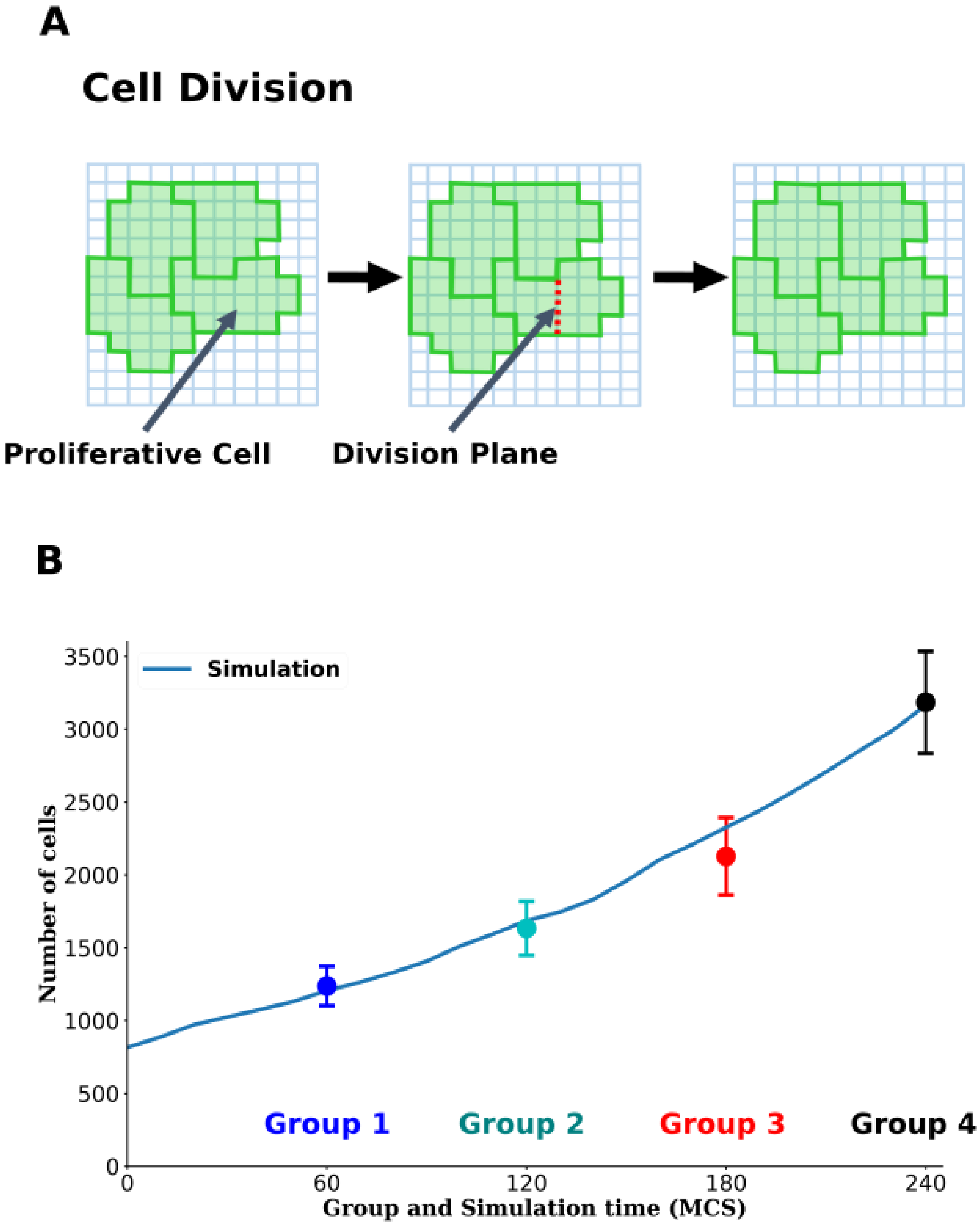


**Figure S8. Cell proliferation in the cellular level of the multi-scale model.**

(A) Schematic representation of cell division in the model. The division plane is defined as the shortest 'diameter' of the proliferative cell. (B) Comparison of the average number of cells within the wing pouch as a function of the group number and the number of cells in the simulations (blue curve). Groups 1, 2, 3, and 4 correspond to 60, 120, 180, and 240 Montecarlo Steps (MCS).

### 7. Simulated Vg data extraction and normalization

To quantitatively extract the experimental and simulated Vg expression space profile, we selected a region of the simulated wing pouch as in the experimental one. First, we selected a rectangular region of the pouch with the same criteria as in experimental wing discs [except that here we only considered the dorsal compartment, and measure the DV distance from the maximum value of Vg to the dorsal-most end of the disc (Figs 2B-E and S9)]. Then, we normalized both Vg intensity and DV distance by their maximum values, and horizontally averaged the Vg concentrations to obtain the normalized Vg concentration as a function of the normalized DV distance binned in 50 intervals as in our experiments.

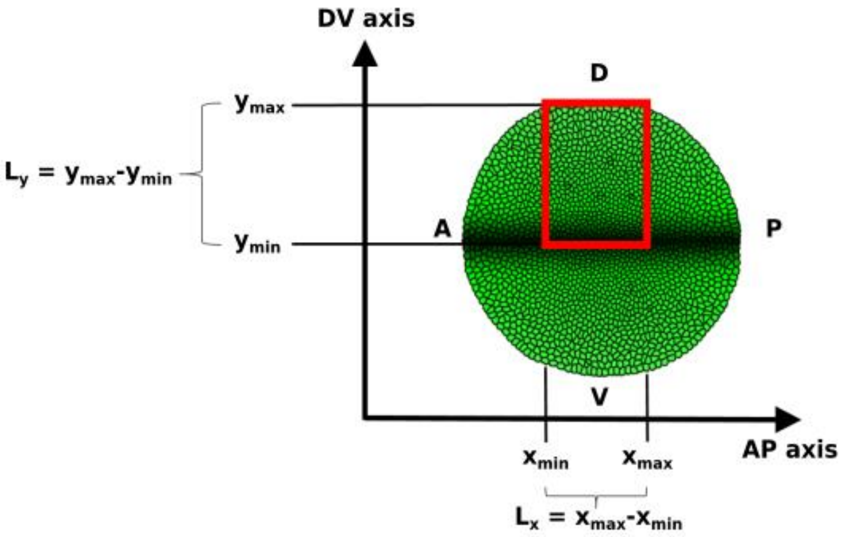


**Figure S9. Selection of the region in simulated discs to compare the model-predicted Vg expression profile with experimental data.**

Selected an area (of $L_y$ times $L_x$, defined by the red rectangle, where $L_y$ and $L_x$ are the lengths along the DV and the anterior-posterior (AP) axis, respectively) of the dorsal compartment of our simulated wing pouch. $L_y$ is the distance between the DV boundary and the dorsal end of the disc whereas $L_x$=0.6 $L_y$ as in experimental discs.

### 8. Fitting score function

We fitted the model to the experimental data by varying the parameters $\alpha$, $\beta$, $\gamma$ and $\delta$, from Eqn. S5 and searching for the minimal value of a Score function $S$, defined as follows:

$$S = \sum_{groups} \sum_{bins} \frac{(Vg_{exp} - Vg_{sim})^2}{\mathrm{sem}_{exp}} \qquad \text{(Eqn. S8)}$$

where $Vg_{exp}$ represents the average experimental Vg intensity in the interval for the corresponding group profile, $\mathrm{sem}_{exp}$ is the standard error of the mean of that data, and $Vg_{sim}$ is the corresponding value in the simulations, the first sum is over the DV binned intervals considered (*i.e.*, 1 to 50) and the second is over the groups compared (*i.e.*, 1 to 4). The best fitting parameters' values for both, the model involving (Fig. 2B"-E", Fig. S10A, and Fig. S11A-D) and lacking recruitment (Fig. 2B-E, Fig. S10B, and Fig. S11A, D) are shown in Table S3.

| Parameter | Model with recruitment (A.U.) | Model without recruitment (A.U.) |
|---|---|---|
| $\alpha$ | 0.085 | 0.12 |
| $\beta$ | 0.191 | 0.0 (imposed) |
| $\gamma$ | 0.5 | - |
| $\delta$ | 0.23 | 0.0 |

**Table S3.** Best fitting values of the Eqn. S5 parameters.

### 9. Comparing models with and without recruitment by the Akaike information criterion

To choose between the models having and lacking recruitment (Fig. 2B"-E", Fig. S10A,B), we applied the second-order Akaike information method (Akaike, 1974). Accordingly, after model dependent fits, we selected the model that provided the lowest Akaike score (AIC). While the best-fitted simulation from the model with recruitment has an AIC of about -17,249, the model without recruitment shows a score of -7,120 and the evidence ratio between both models diverges. Altogether, this analysis indicates that the model with recruitment has more relative likelihood than the model lacking recruitment to reproduce the here reported experimental data.

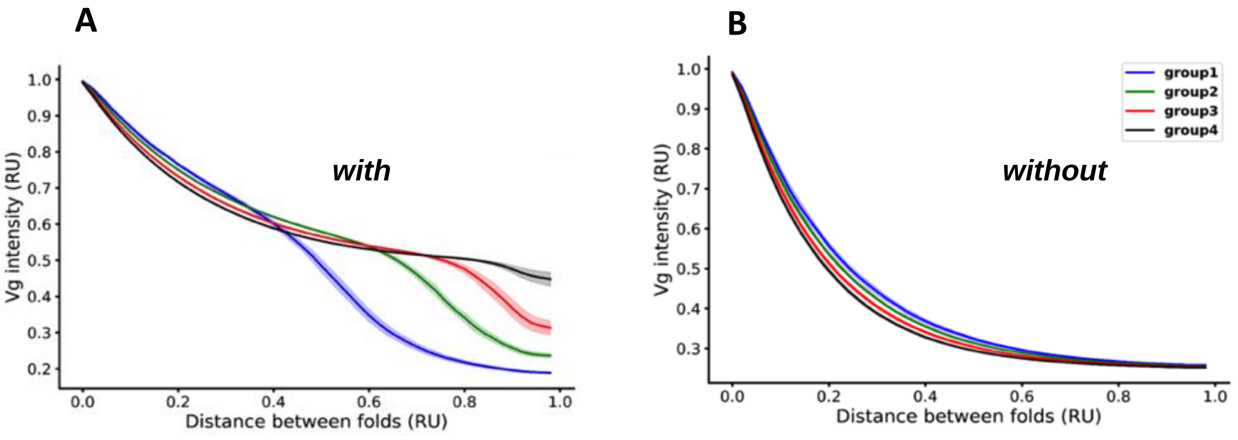


**Figure S10. While the presence of a recruitment mechanism produces over-scaling, its absence produces a slight under-scaling.**

The normalized profiles obtained for best fitting parameters for both, the model involving (A) and lacking (B) recruitment. The profiles are colored as follows: group 1 in blue, group 2 in green, group 3 in red and group 4 in black, for both models. In each case, the curves represent the mean values over 10 simulations, and the colored shadows are the standard deviations.

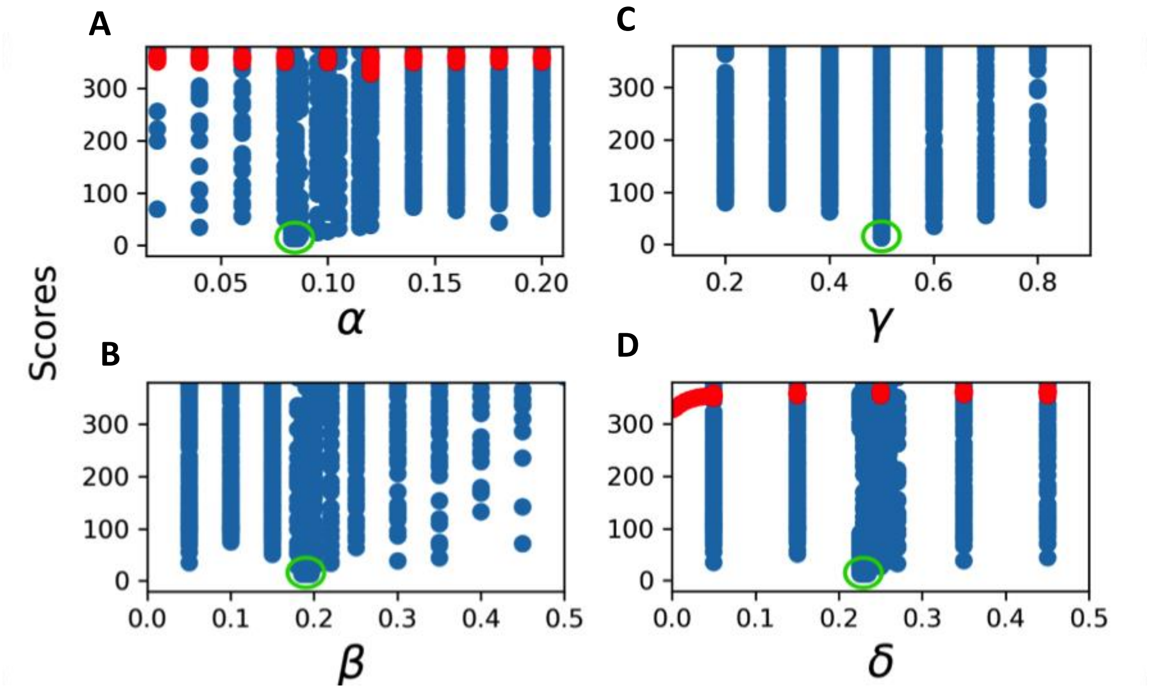


**Figure S11. Parameters space exploration showing that a model with recruitment has lower scores than the model without recruitment.** Scores obtained by sweeping the parameters $\alpha$, $\beta$, $\gamma$ and $\delta$ for the model with recruitment (blue dots) and $\alpha$ and $\delta$ for the model without recruitment (red dots). The best-fit parameter values are marked by green circles.

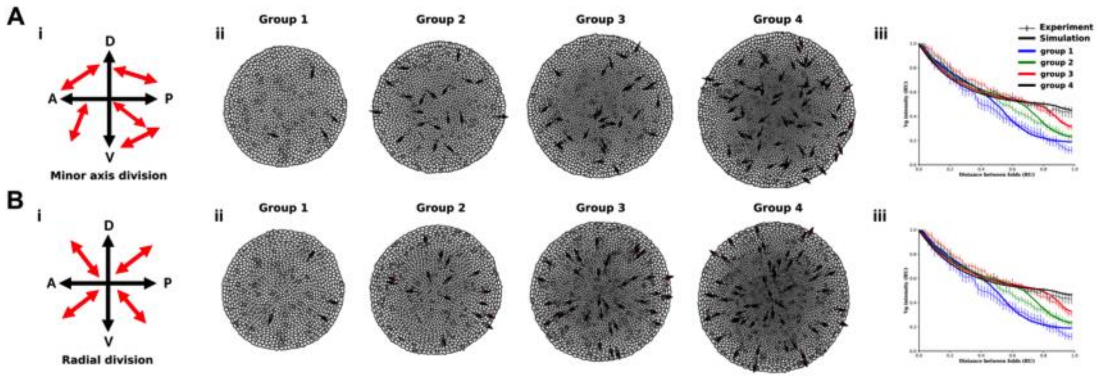


**Figure S12 - Orientation of cell division does not affect cell recruitment**. The mathematical model presented in Fig. 2 was also simulated in a condition in which cell divisions are oriented in a radial direction rather than along the minor axis of the cell (red arrows). Cell division is along the minor axis (A), or in a radial direction from the center of the simulated tissue (B). In each condition, from left to right, it can be observed (i) the direction of cell division (red arrow), (ii) the simulated tissues for groups 1 to 4 (in this representation, cells about to divide are marked with an arrow in the direction of division), and (iii) the corresponding model-predicted Vestigial profiles of the four groups overlapped with the experimental data. Model parametrization is as in Fig. 2.

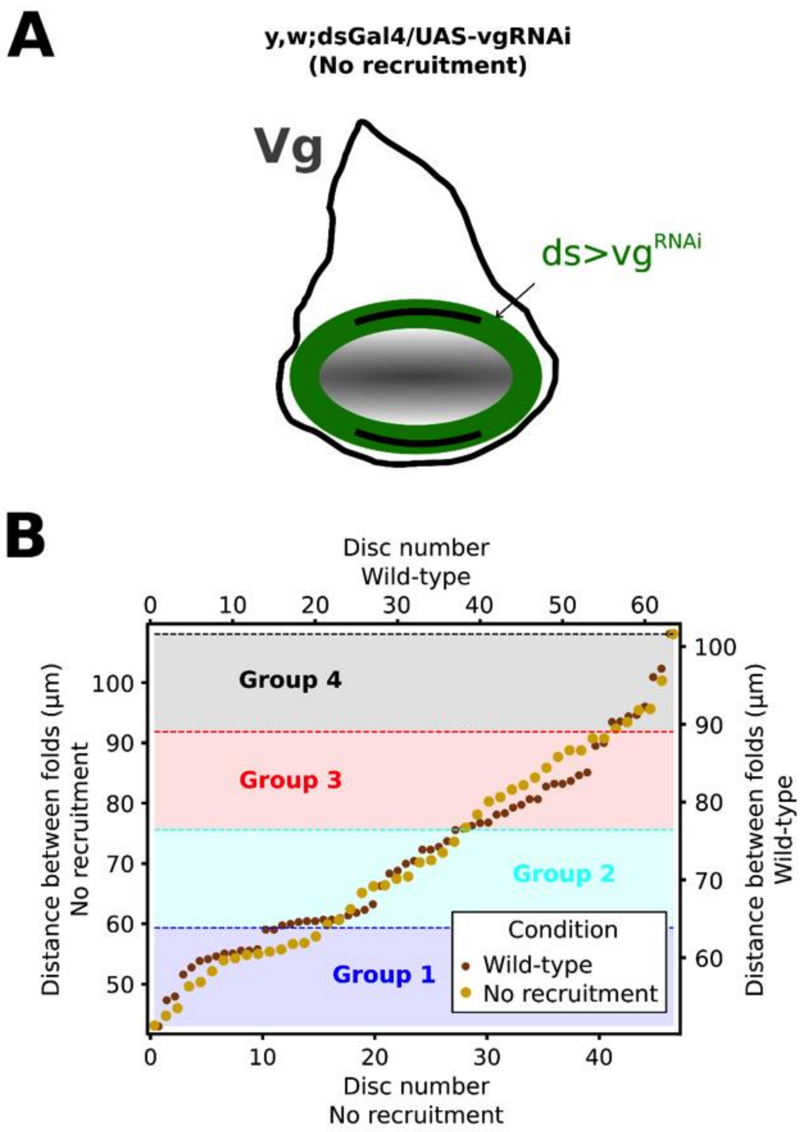


**Figure S13. Schematics of recruitment inhibition and comparison of discs' sizes**.

(A) Recruitment is inhibited by expressing $vg^{RNAi}$ in the pattern of *ds* (green domain) using the Drosophila Gal4-UAS system (denoted by $ds{>}vg^{RNAi}$). (B) Distribution of *y*, *w* (wild-type condition) and *y*, *w*; dsGal4 / UAS-$vg^{RNAi}$ (recruitment-impaired) disc sizes according to their DV length and their subdivision into four groups. (As in Fig. 1B, groups are determined by dividing the interval defined by the smallest and largest DV length into 4 equal parts).

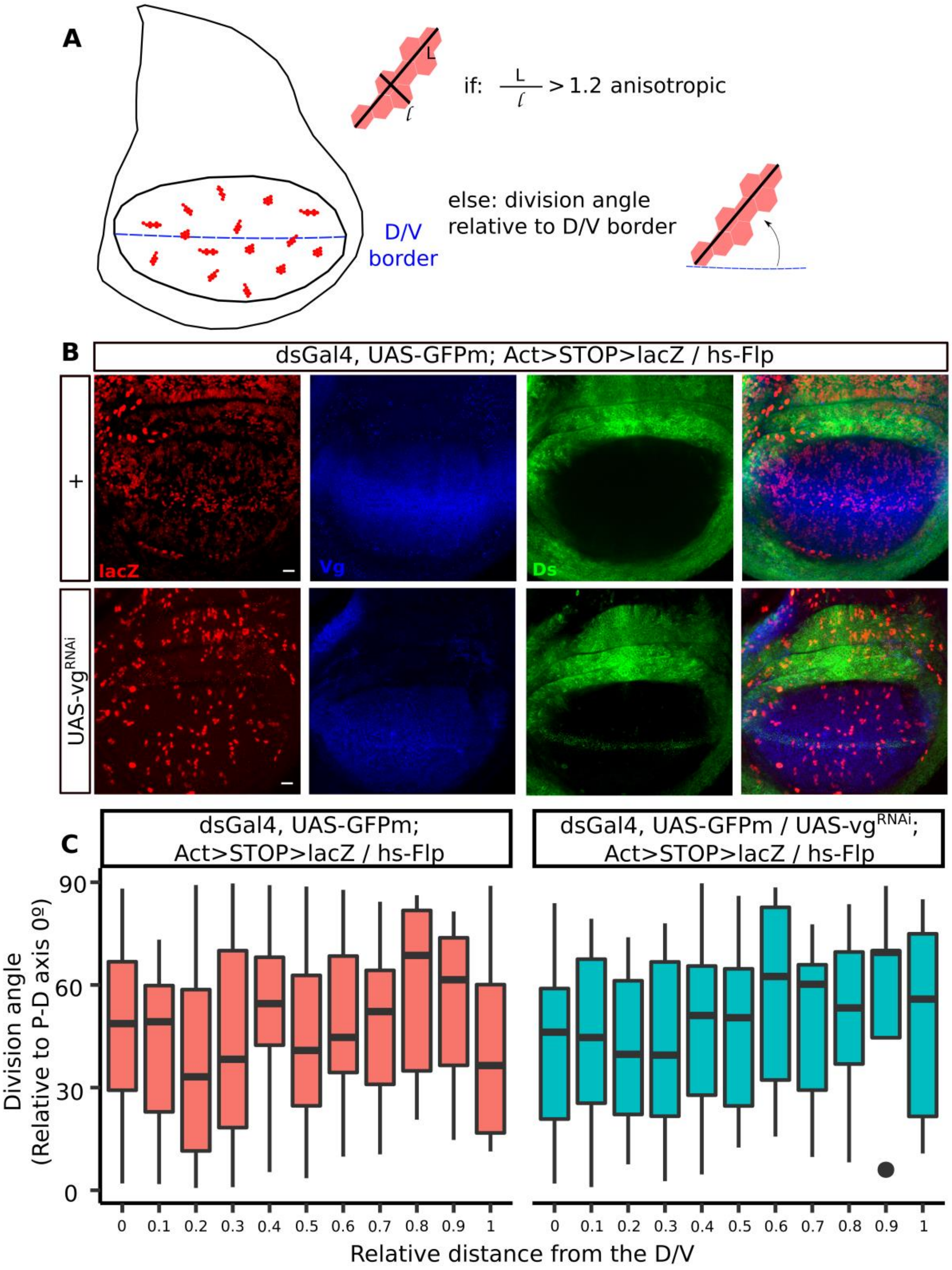


**Figure S14. Orientation of cell division is similar in control and *ds>vg*$^{RNAi}$ .**

Clone orientations and division orientations change along the P–D axis. (A) Clones marked by expression of lacZ were induced at early third instar by a 10-min heat shock at 38ºC to induce in cis FLP/FRT- mediated recombination. Diagram representing an imaginal wing disc with red clones within the pouch, major (L) and minor (l) axis for each clone and the ratio was determinate. The long axis of each dividing clone is oriented relative to the P–D axis and plotted against its relative distance from the center to the first fold (edge) of the pouch (C).(B) Third instar imaginal wing discs containing clones expressing lacZ. (C) Box plots show median and first and third quartiles. Only cells with elongation ratios (long/short axis) >1.2 are plotted.

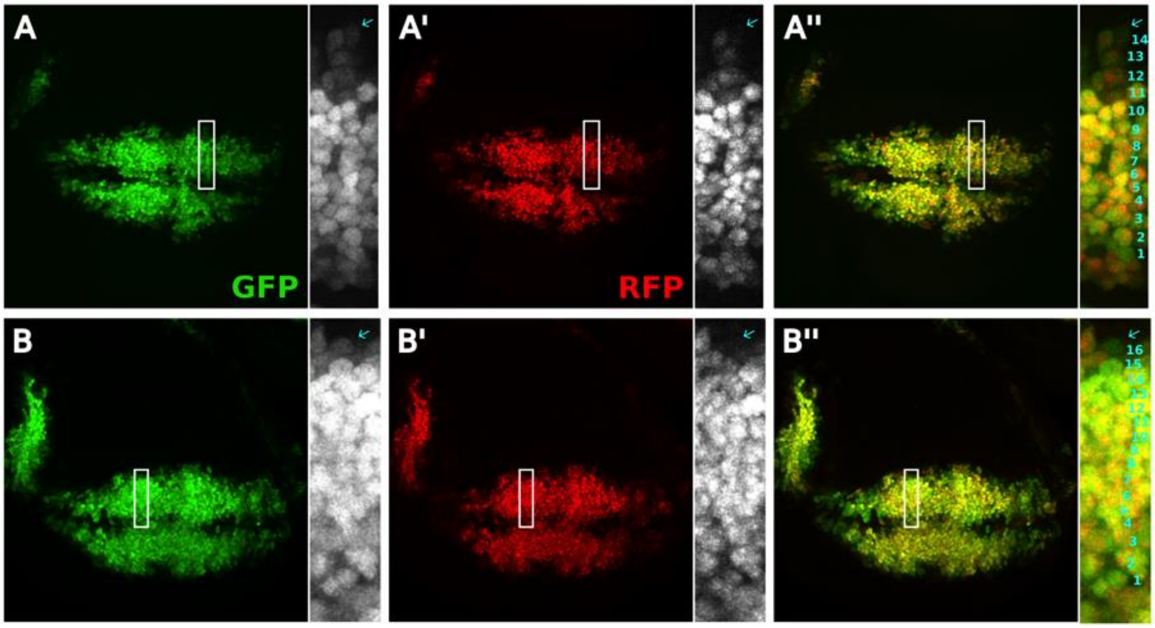


**Figure S15. Vg is established beyond the region of Wg signaling**.

Immunofluorescence images of two representative *vg*$^{QE}$Gal4 UAS-TransTimer wing discs with DV length in the 60 – 80 μm range. Images show the expression of GFP (A, B), RFP (A', B'), or both (A'', B''). At the right of each panel, an amplification of the region inside the white rectangle is shown and cells from the DV border are counted as in Chaudhary *et al.* 2019. The cyan arrow points to newly-recruited cells (GFP positive, RFP negative) located 14 (A) or 16 (B) cells away.
